# Developmental conversion of the nucleolus into an RNA Polymerase II transcriptional platform in *Drosophila* spermatocytes

**DOI:** 10.64898/2026.05.20.726666

**Authors:** Jaclyn M. Fingerhut, Jun I. Park, Rebecca Y. Li, Romain Lannes, Archana Ashok, Yukiko M. Yamashita

## Abstract

The nucleolus is widely regarded as a specialized compartment for RNA polymerase I (Pol I)-driven ribosomal RNA transcription and ribosome biogenesis. Yet the presence of “atypical nucleoli”, or nucleolus-like bodies (NLBs), which lack rRNA transcription despite containing canonical nucleolar components, has long been recognized, most notably during mammalian oogenesis and spermatogenesis. NLBs have been shown to have an essential function independent of rRNA transcription, but the nature of that function remained unclear. Here, we demonstrate that the nucleolus becomes an NLB during spermatocyte development in *Drosophila melanogaster* and, surprisingly, that this NLB serves as a platform for RNA polymerase II (Pol II)-mediated transcription. We find that the Y chromosome-linked fertility genes, which are heterochromatic in most cell types but highly expressed in spermatocytes, are transcribed at the spermatocyte NLB. We further show that the recruitment of active Pol II to the NLB requires known spermatocyte-specific transcriptional regulators. In their absence, the Y-linked fertility genes embedded within heterochromatin are not properly transcribed. Our findings reveal an active role for an NLB as a Pol II platform, and we propose that other NLBs may have similar functionality.

## Introduction

The nucleolus is commonly viewed as a nuclear compartment for RNA polymerase I (Pol I)-mediated ribosomal RNA (rRNA) transcription and ribosome assembly^1, 2^. However, the nucleolus was originally recognized cytologically as a prominent, phase-dark structure within the nucleus long before its molecular basis was understood. The nucleolus was subsequently linked to specific chromosomal loci, termed nucleolar organizer regions (NORs), which were later shown to correspond to rDNA loci, the repetitive gene cluster encoding rRNAs. Further studies established the nucleolus as the hub of Pol I-mediated rRNA transcription and ribosome biogenesis. Recent studies have led to the understanding that the nucleolus is a biomolecular condensate whose multilayered phase-separated organization is thought to underlie compartmentalization of the ribosome biogenesis processes, supporting step-wise ribosome maturation^3, 4^. Together, these discoveries have shaped the view of the nucleolus as a Pol I platform.

However, interestingly, the presence of “atypical nucleoli” lacking active rRNA transcription and ribosome biogenesis has long been recognized in mammalian spermatids, oocytes, and early embryos^5–7^. In mammalian oocytes, atypical nucleoli, termed nucleolus-like bodies (NLBs), are formed from the canonical nucleolus as rRNA synthesis ceases over the course of oocyte maturation^8^. NLBs are negative for rRNA transcription and lack the typical tripartite structure of the nucleolus^9–11^. However, NLBs still contain canonical nucleolar proteins (such as Fibrillarin and Nucleolin) and are surrounded by heterochromatic DNA^5, 6, 11^. Upon fertilization, NLBs become known as nucleolar precursor bodies (NPBs), reflecting the initial assumption that they serve as a reservoir for reconstituting rRNA-transcribing nucleoli in the developing embryo^5, 6, 10, 12^. However, removal of the NPB did not prevent the activation of rRNA expression^13^, yet did result in embryonic lethality^14, 15^, suggesting that NLB/NPB’s essential function is independent of rRNA transcription. The nature of that function remains a mystery.

*Drosophila* spermatogenesis has served as an excellent model system to study cellular differentiation. Upon completion of the mitotic divisions, spermatocytes enter an extended meiotic G2 phase, lasting 80-90 hours, during which an extensive spermatocyte-specific gene expression program expresses a suite of genes required for meiosis and sperm development (Fig 1A)^16, 17^. This specialized gene expression program depends on two groups of transcriptional regulators: the testis-specific TBP-associated factors (tTAFs) and the testis-specific meiotic arrest complex (tMAC)^18^. Intriguingly, the tTAFs have previously been shown to localize to the nucleolus^19, 20^. However, the nucleolus was presumed to lack a direct role in RNA polymerase II (Pol II)-mediated transcription. Therefore, localization of the tTAFs to the nucleolus was proposed to have an indirect role in spermatocyte gene expression through sequestering Polycomb repressive complexes into the nucleolus, thereby relieving repression at target loci elsewhere in the genome^19^.

**Figure 1:**
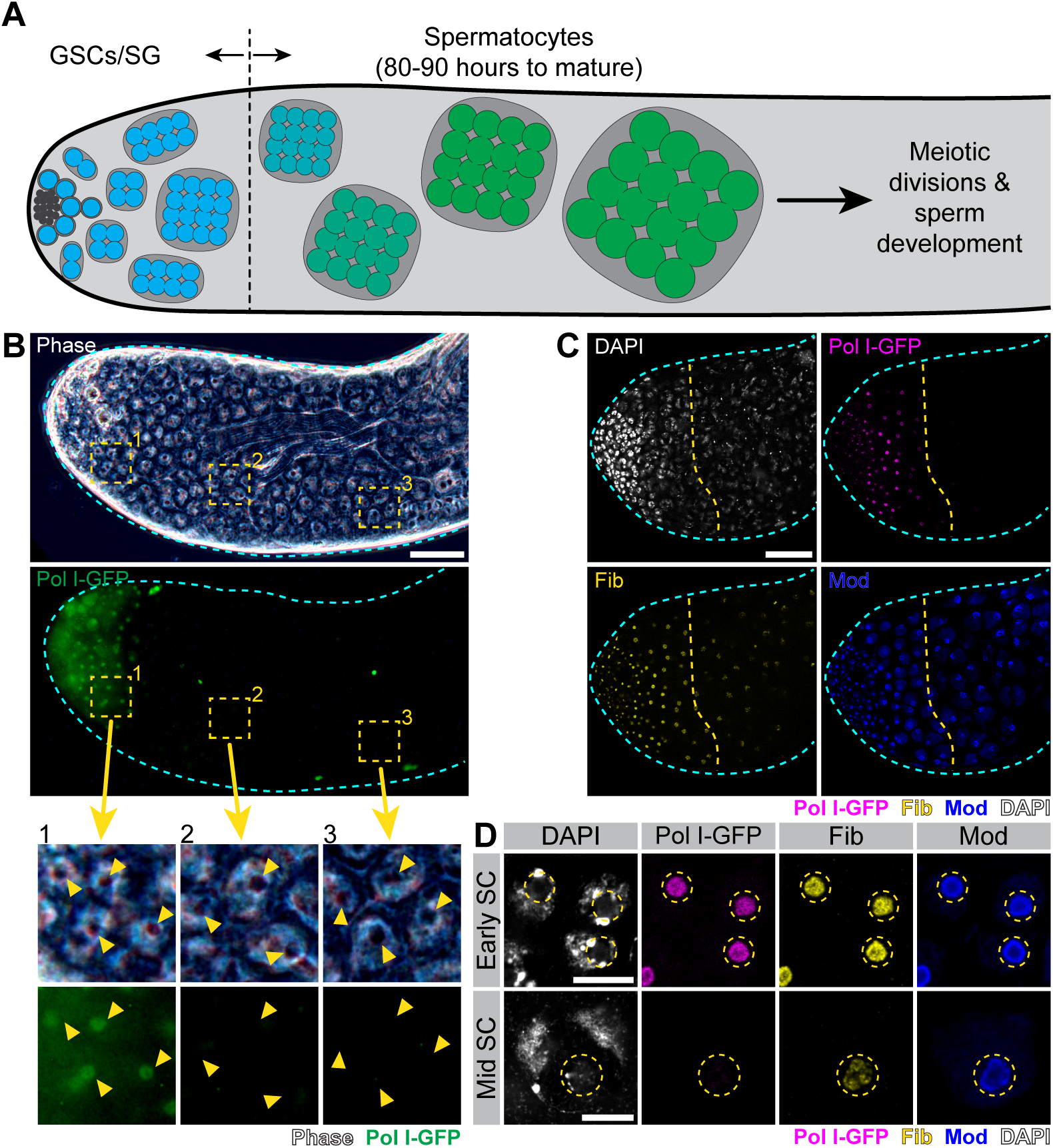
The spermatocyte nucleolus converts into an NLB (A) Diagram of *Drosophila* germ cell development through the end of spermatocyte maturation. Germline stem cells (GSCs, blue) localize around somatic niche cells (dark gray) at the apical tip. Following GSC division, germ cells are known as spermatogonia (SG, blue) and undergo several rounds of mitosis before becoming spermatocytes (green). Germ cells develop as cysts encapsulated by somatic cyst cells (gray). (B) Top: Phase contrast image of the apical tip of a testis (cyan dashed line) through the end of spermatocyte development also expressing Pol I-GFP (Polr1B-GFP, green). Bar: 50µm. Bottom: Enlarged images of the three regions in the yellow dashed squares above. Yellow arrowheads mark nucleoli/NLBs. (C) Immunofluorescence staining of Fib (yellow), Mod (blue), and DAPI (white) in the apical tip of a testis (cyan dashed line) through mid-spermatocytes also expressing Polr1B-GFP (magenta). The yellow dashed line indicates the developmental point at while Pol I no longer occupies the nucleolus/NLB. Bar: 50µm. (D) Immunofluorescence staining of Fib (yellow), Mod (blue) and DAPI (white) in early and mid spermatocytes (SCs) also expressing Polr1B-GFP (magenta). Yellow dashed lines encircle nucleoli/NLBs. Bar: 5µm.

Among the most striking examples of genes expressed in spermatocytes are the Y chromosome-linked fertility genes. The *Drosophila* Y chromosome has only 12 single-copy protein-coding genes, many of which are essential for male fertility^21–23^. The Y-linked fertility genes have several unique characteristics. First, the majority of the Y-linked fertility genes are heterochromatic in other cell types but are robustly transcribed in spermatocytes^24–26^. This robust transcription is associated with a so-called ‘lampbrush chromosome’, where many Pol II enzymes are loaded onto the DNA to facilitate high levels of transcription^27^. Second, many of the Y-linked fertility genes contain long stretches of satellite DNA (non-coding repetitive DNA) in their introns, resulting in genes that exceed megabases in size, a phenomenon known as intron gigantism^26, 28–30^. These intronic satellite DNAs, as well as intergenic satellite DNAs and transposons distributed throughout the Y chromosome, contribute to the heterochromatic nature of the Y-linked genes. How the heterochromatic Y chromosome becomes compatible with such robust transcription in spermatocytes remains poorly understood.

Here, we show that the nucleolus becomes an NLB during spermatocyte development in *Drosophila*, converting from a canonical nucleolus in spermatocyte progenitors to an NLB lacking Pol I and rRNA in spermatocytes. Surprisingly, we find that the spermatocyte NLB becomes associated with Pol II and serves as a Pol II platform, facilitating the expression of Y-linked fertility genes. We find that the tTAFs and tMAC are necessary for the recruitment and/or activation of Pol II at the NLB to transcribe the Y-linked fertility genes. We further show that tMAC promotes the opening of these heterochromatic Y-linked loci. These results suggest that NLB-localized tTAFs and tMAC may have a direct role in Pol II-mediated transcription. Together, our findings reveal that the NLB of *Drosophila* spermatocytes functions as a Pol II platform to support germ cell development, raising the possibility that other NLBs may also function as Pol II platforms, instead of storage bodies as suggested previously^6, 31^. More broadly, we propose that the nucleolus/NLB may provide a specialized environment for aspects of transcription that are shared between rDNA and the Y-linked fertility genes: transcription of heterochromatic loci and/or robust gene expression, requiring the loading of many polymerases beyond the normal constraints of transcriptional regulation.

## Results

### Nucleoli in developing *Drosophila* spermatocytes convert into NLBs

The extensive literature on *Drosophila* spermatocytes has long recognized a prominent ‘nucleolar’ structure, identifiable by phase microscopy (Fig 1B), which is also positive for canonical nucleolar proteins such as Fibrillarin (Fib) and Modulo (Mod, the *Drosophila* homolog of Nucleolin^32^) (Fig 1C, D) ^33, 34^. To our surprise, we found that these nucleoli become negative for Pol I (visualized with Polr1B-GFP) as spermatocyte differentiation progresses (Fig 1B-D).

In mouse spermatogenesis, it has been shown that the nucleolus also ceases rRNA transcription, transforming into an atypical nucleolus that still contains other canonical nucleolar proteins ^7, 35^. This led us to hypothesize that the nucleolus in *Drosophila* spermatocytes may also represent an NLB. Indeed, we found that nucleolar rRNA decreased in spermatocytes, coinciding with the disappearance of Pol I (Fig 2). Pol I transcribes rDNA from a single promoter in the ETS, generating a single cistronic transcript that is then cleaved to yield the 18S, 5.8S and 28S rRNAs (Fig 2A)^36^. RNA fluorescent *in situ* hybridization (FISH) showed that the nucleolar 18S and 28S rRNAs became markedly reduced as Pol I signal diminished from the nucleolus (Fig 2B). Moreover, unprocessed rRNA (detected by probes that hybridize to the junctions between ITS1 and 18S or ITS2 and 28S (Fig 2A)), which better mirrors ongoing rRNA transcription^37^, also exhibited a marked decrease over spermatocyte differentiation (Fig 2C), confirming that rRNA transcription mostly ceases during spermatocyte development. Importantly, although rRNA transcription decreases during spermatocyte development, enough ribosomes are stored in the cytoplasm to allow for sperm development, because cytoplasmic rRNA was abundantly observed throughout the rest of germ cell development (Fig S1).

**Figure 2:**
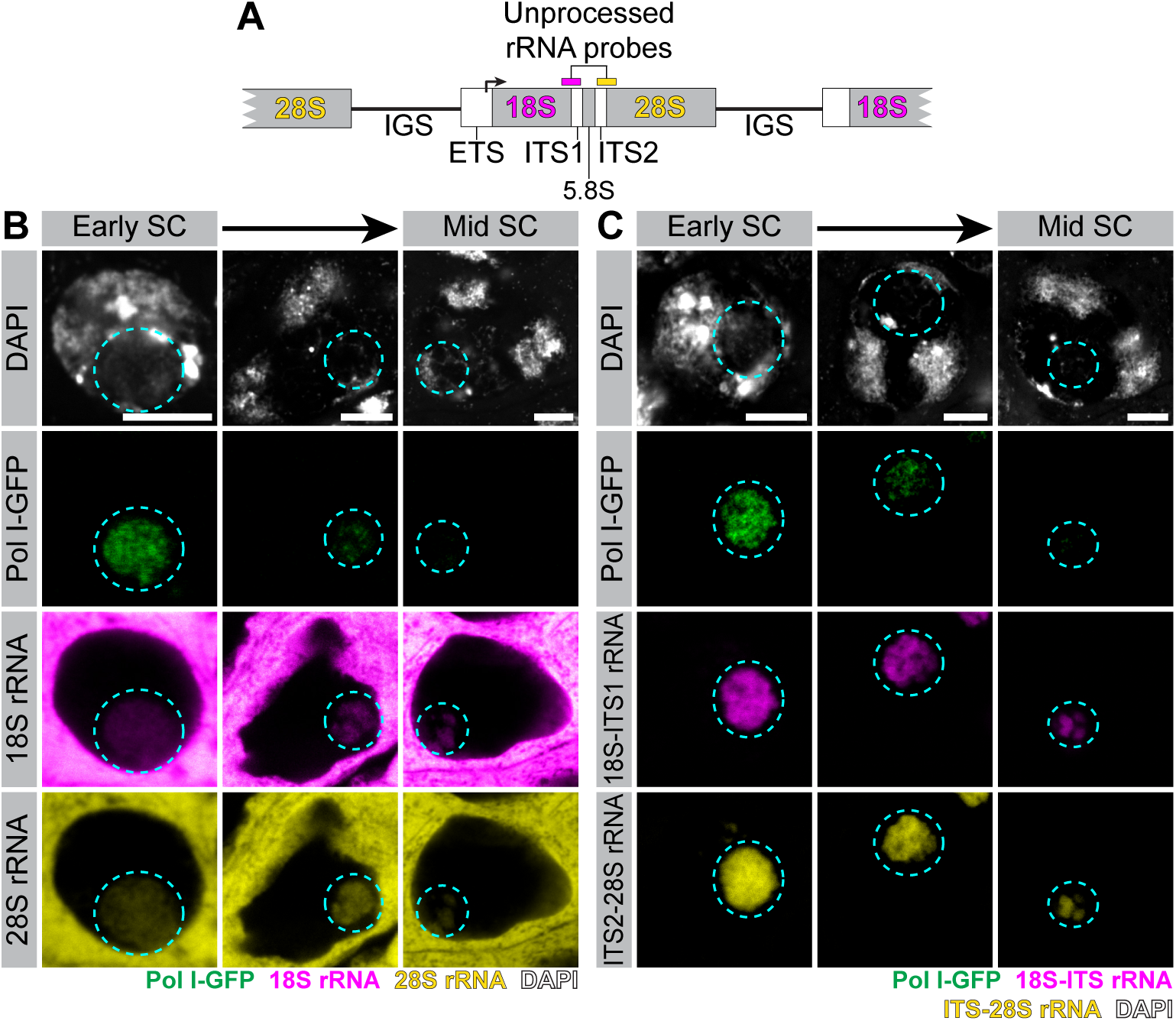
rRNA transcription diminishes during spermatocyte development (A) Diagram of an rDNA cistron showing the rRNA genes (18S, 28S, 5.8S) and spacer sequences (IGS, ITS1, ITS2) as well as the transcription start site in the ETS. Locations of unprocessed rRNA probes are indicated by bars above the diagram. (B) RNA FISH for 18S (magenta) and 28S (yellow) rRNAs in spermatocytes of different stages also expressing Polr1B-GFP (green) and stained with DAPI (white). Nucleolus/NLB encircled with cyan dashed line. Bars: 5µm. (C) RNA FISH for unprocessed 18S-ITS1 (magenta) and ITS2-28S (yellow) rRNAs in wildtype spermatocytes of different stages also expressing Polr1B-GFP (green) and stained with DAPI (white). Nucleolus/NLB encircled with cyan dashed line. Bars: 5µm.

Together, these results demonstrate that Pol I-mediated transcription of rDNA largely ceases during spermatocyte development, whereas the nucleolar structure itself (as seen by Fib and Mod staining) remains largely intact. We therefore conclude that the nucleolus in *Drosophila* spermatocytes transitions into an NLB.

### Pol II localizes to the NLB during spermatocyte development

Given the absence of Pol I and rRNA transcription, NLBs have long puzzled researchers. Surprisingly, we found that Pol II became concentrated at the spermatocyte NLB, coinciding with the disappearance of Pol I (Fig 3A, B, Fig S2A). In mitotically dividing germ cells (stem cells and spermatogonia) and very early spermatocytes, Pol II mainly overlapped with DAPI-stained DNA, consistent with its known role in transcription (Fig 3B, Fig S2A). However, in maturing spermatocytes, Pol II strongly concentrated as a ring surrounding the NLB (Fig 3 A - C). Multiple methods of Pol II detection [anti-Pol II staining (Fig 3A, B) and endogenously tagged Pol II subunits (Polr2C-GFP and mCherry-Polr2A) (Fig S2B, C)] consistently showed Pol II localization at the NLB, confirming that Pol II is indeed recruited to the NLB. Moreover, initiating Pol II-specific phosphorylation (pSer5, phosphorylated at Serine 5 in the C-terminal domain (CTD)), and elongating Pol II-specific phosphorylation (pSer2, phosphorylated at Serine 2 in the CTD)^38^, also exhibited localization to the NLB (Fig 3D, E), indicating that NLB-localized Pol II is actively engaged in transcription.

**Figure 3:**
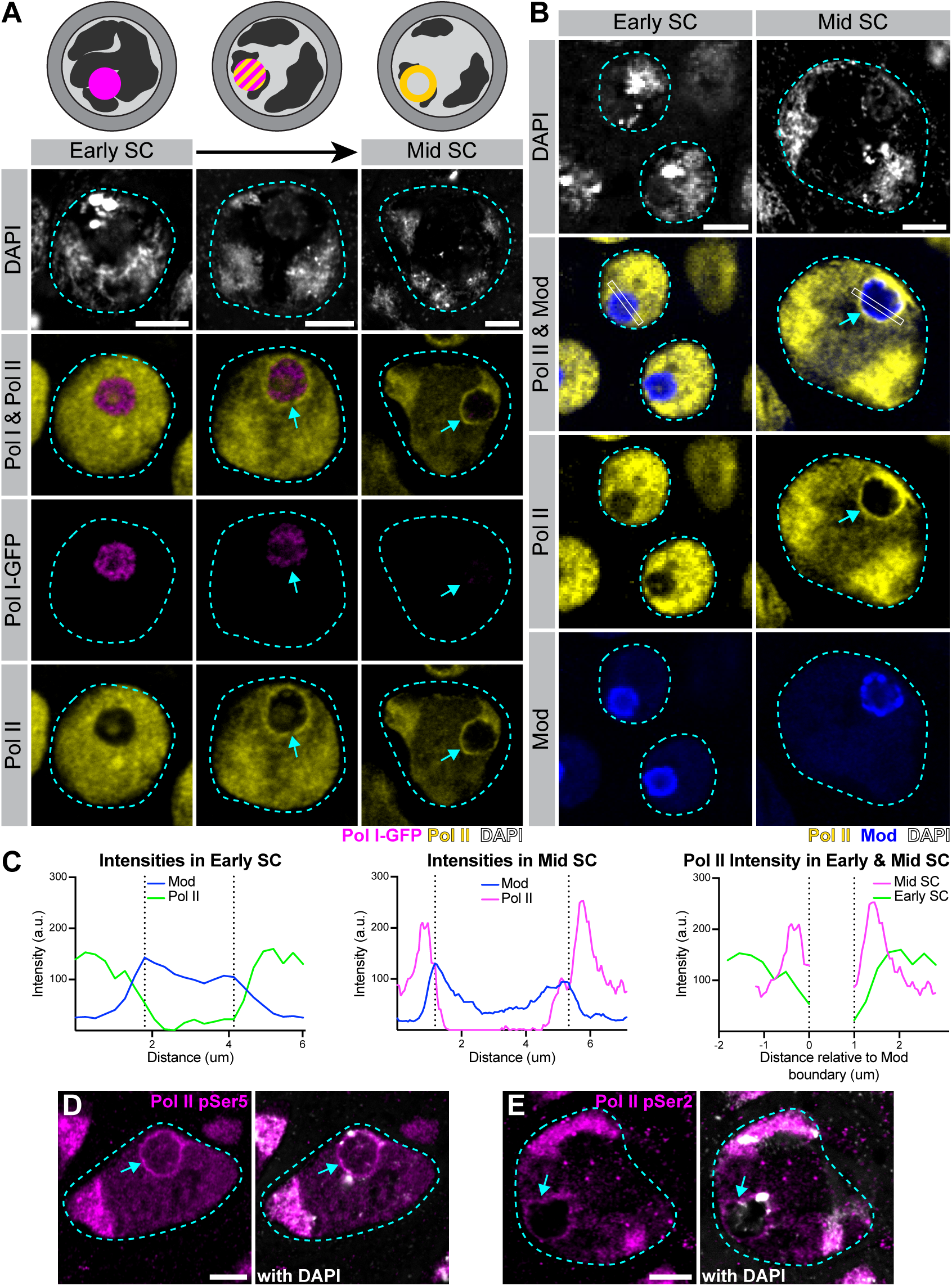
Pol II localizes to the spermatocyte NLB (A) Top: Diagrams of spermatocytes illustrating the conversion from Pol I (magenta) to Pol II (yellow) in the nucleolus/NLB. Bottom: Immunofluorescence staining of Pol II (yellow), and DAPI (white) in spermatocytes (cyan dashed line) of different stages also expressing Polr1B-GFP (magenta). Nucleolar/NLB-localized Pol II indicated by cyan arrows. Bars: 5µm. (B) Immunofluorescence staining of Pol II (yellow) and Mod (blue) and DAPI (white) in early or mid spermatocytes. Nuclei outlined with cyan dashed line, nucleolar/NLB-localized Pol II indicated with a cyan arrow. Bars: 5µm. (C) Intensity plots for Pol II (green or magenta) and Mod (blue) for the regions in white rectangles in (B) (left, center). The plot on the right equates the nucleolar/NLB boundary defined by Mod signal drop off in the left and center plots (vertical dashed lines) as 0 or 1 allowing for a comparison of Pol II intensity in early (green) and mid (magenta) spermatocytes relative to Mod. (D) Pol II pSer5 (magenta) expression in a mid spermatocyte NLB (cyan dashed outline) with DAPI (white). NLB marked by cyan arrow. Bar: 5µm. (E) Pol II pSer2 (magenta) expression in a mid spermatocyte NLB (cyan dashed outline) with DAPI (white). NLB marked by cyan arrow. Bar: 5µm.

Together, these results demonstrate that the spermatocyte NLB contains active Pol II. The prominent localization of Pol II within the spermatocyte NLB is particularly striking as Pol II localization to the NLB in mammalian oocytes has been debated in the literature^6, 31^. Our results also demonstrate that the transition from Pol I to Pol II in the nucleolus/NLB is a continuous process occurring over early-to-mid spermatocyte development.

### The Y-linked fertility genes are transcribed at the spermatocyte NLB

Given that rRNA transcription ceases coinciding with Pol II recruitment to the NLB, the transcripts produced by NLB-associated Pol II are unlikely to be rRNA. Interestingly, the presence of RNA (which was shown not to be rRNA) has been noted in mouse NLBs^6, 10, 11^. If not rRNA, what are the transcriptional products of NLB-associated Pol II in spermatocytes?

Surprisingly, we found that localization of Pol II to the NLB coincides with the burst of transcription associated with the Y-linked fertility genes, such as *kl-3, kl-5* and *ORY* (Fig 4A-C, Fig S3). While the Y-linked fertility genes are not transcribed in mitotic cells, they are required for male fertility and become highly transcribed in developing spermatocytes^22, 23, 26, 28^. Due to their large size owing to gigantic introns, transcription of these genes takes the entirety of spermatocyte development, spanning ∼80-90 hours^39^. The earlier exons of *kl-3* are first expressed in very early spermatocytes, followed by expression of the satellite DNA-containing large introns in more differentiated spermatocytes (Fig 4A). Subsequently, mature mRNA becomes detectable only in the cytoplasm of very mature spermatocytes in the form of RNP granules^39, 40^. Interestingly, we noted that the recruitment of Pol II to the NLB became most prominent as the burst of intronic transcripts became detectable, whereas the transcription of the first exon was observed before the clear recruitment of Pol II to the nucleolus (Fig 4B, C). A similar pattern was observed for the Y-linked gigantic genes *kl-5* and *ORY* (Fig S3). These results suggest that the spermatocyte NLB may play an important role in achieving robust transcription of the Y-linked fertility genes, although initiation may occur without the NLB.

**Figure 4:**
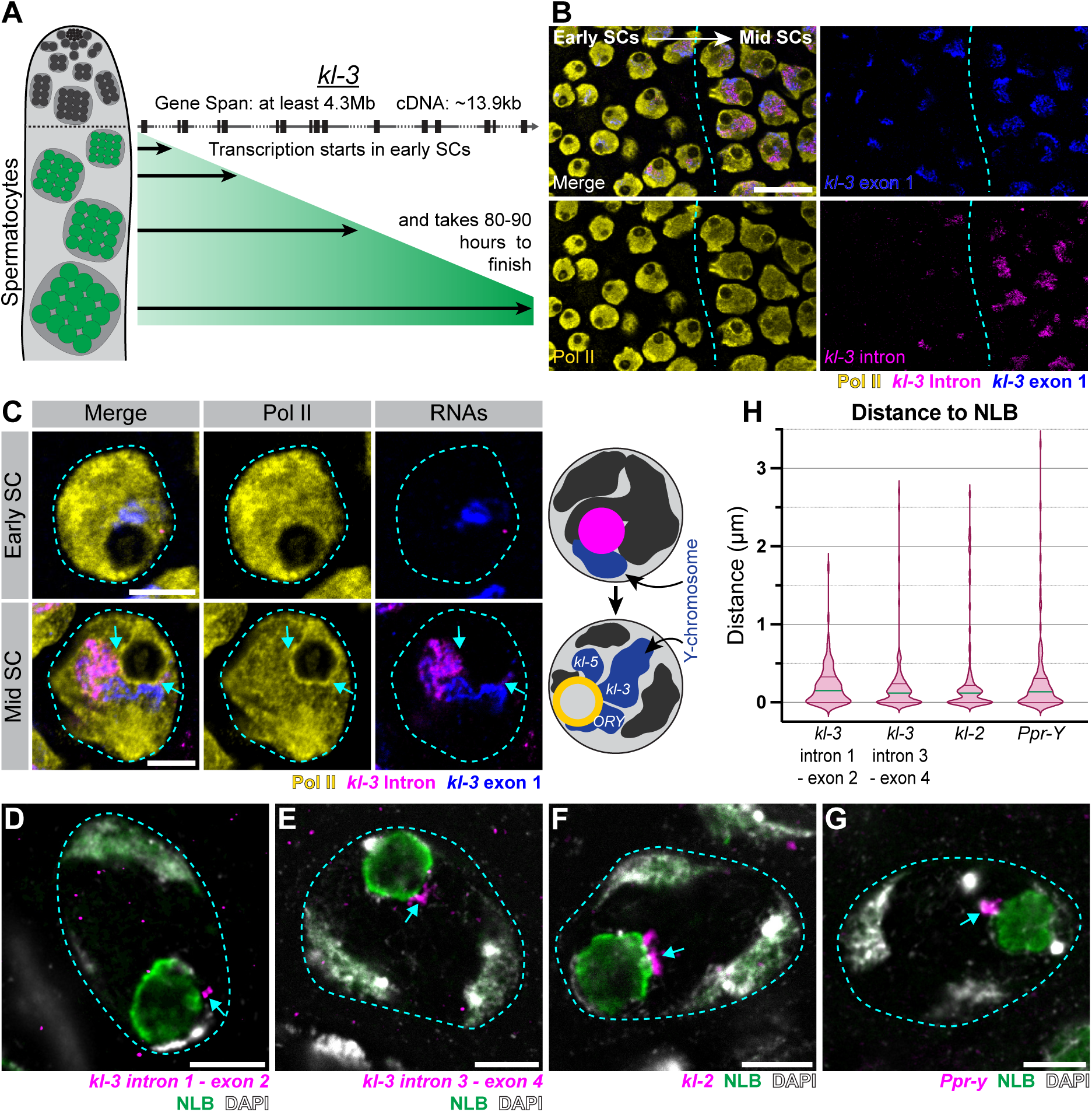
The Y-linked fertility genes are transcribed at the spermatocyte NLB (A) Diagram of *kl-3* transcription over spermatocyte development. Left: Spermatocytes are in green while other germ cells and somatic cells of the testis are gray scale. Right: The top diagram shows the *kl-3* gene with exons as vertical rectangles and introns containing satellite DNA repeats as dashes. Below, arrows indicate how much of *kl-3* is expressed in the corresponding spermatocyte developmental stage on the left. (B) Immunofluorescence staining for Pol II (yellow) with RNA FISH for the first exon (blue) and intronic repeats (magenta) of the Y-linked genes *kl-3* in early-to-mid spermatocytes. Vertical cyan dashed line indicates the point at which Pol II becomes nucleolar. Bar: 20µm. (C) Left: Immunofluorescence staining for Pol II (yellow) with RNA FISH for the first exon (blue) and intronic repeats (magenta) of the Y-linked genes *kl-3* in early and mid spermatocytes. Spermatocyte nuclei are outlined with cyan dashed lines. Cyan arrows mark nucleolar/NLB-localized Pol II. Bars: 5µm. Right: diagrams depicting how the Y chromosome looping occurs during spermatocyte development and how the Y chromosome fills the nucleoplasm alongside the switch from Pol I to Pol II in the nucleolus/NLB. (D) HCR RNA FISH for the *kl-3* intron 1 – exon 2 junction in a mid spermatocyte expressing Sa-GFP, which marks the NLB. Spermatocyte nucleus outlined with cyan dashed line and RNA signal indicated by a cyan arrow. Bar: 5µm. (E) HCR RNA FISH for the *kl-3* intron 3 – exon 4 junction in a mid spermatocyte expressing Sa-GFP. Spermatocyte nucleus outlined with cyan dashed line and RNA signal indicated by a cyan arrow. Bar: 5µm. (F) HCR RNA FISH for the *kl-2* in a mid spermatocyte expressing Sa-GFP. Spermatocyte nucleus outlined with cyan dashed line and RNA signal indicated by a cyan arrow. Bar: 5µm. (G) RNA FISH for the *Ppr-Y* in a mid spermatocyte expressing Sa-GFP. Spermatocyte nucleus outlined with cyan dashed line and RNA signal indicated by a cyan arrow. Bar: 5µm. (H) Graph of the distance between the indicated Y-linked transcripts and the NLB. 100 RNA foci were measured per transcript. The median distances (green line) for the transcripts are 0.15µm, 0.12µm, 0.12µm, and 0.135µm respectively and quartiles are shown by the horizontal dark pink lines.

Indeed, RNA FISH of Y-linked fertility genes showed their close association to the spermatocyte NLB. When *kl-3*’s exon 1 signal first became detectable in early spermatocytes, the transcript was consistently near the nucleolus (Fig 4C), although it was before the localization of Pol II at the nucleolus became prominent. This result suggested that transcription may be initiated at the nucleolus prior to its conversion into an NLB. After Pol II’s nucleolar/NLB localization became clear in later spermatocytes, *kl-3*’s exon 1 signal expanded (Fig 4C), reflecting the collection of many *kl-3* transcripts attached to the template DNA in a lampbrush chromosome structure^26, 39^, precluding the detection of ongoing transcription with this method. However, the use of RNA FISH probes specific to intron-exon junctions^41^ (regions of the nascent transcript that are highly transient) revealed that ongoing transcription of *kl-3* was also localized at the NLB in later spermatocytes (Fig 4D, E).

For other Y-linked fertility genes, such as *kl-2* and *Ppr-Y,* whose RNA signal does not expand as much (presumably because of their relatively smaller gene size compared to *kl-3*), transcripts were also localized consistently near the NLB (Fig 4F, G). The consistent proximity of these transcripts/nascent RNAs to the spermatocyte NLB (Fig 4H) indicates that the Y-linked fertility genes are likely transcribed at the NLB by NLB-localized Pol II, further suggesting active involvement of NLBs in transcription.

### Known spermatocyte-specific transcriptional regulators mediate Pol II recruitment to/activation at the NLB

The unexpected shift from Pol I to Pol II in the nucleolus/NLB of developing spermatocytes prompted the question of how this shift is regulated. Likewise, how is transcription of the Y-linked fertility genes (likely mediated by nucleolar Pol II) regulated? Extensive studies have documented the localization of spermatocyte-specific Pol II regulators, the testis-specific TAFs (tTAFs), to the nucleolus of developing spermatocytes^19, 20^. The tTAFs are paralogs of the canonical TAFs and are also believed to function in the TFIID transcription initiation complex^42, 43^. The testis-specific meiotic arrest complex (tMAC) is a testis-specific version of the MuvB complex (MMB/DREAM) that also regulates spermatocyte-specific gene expression along with the tTAFs^17, 44^. tMAC is believed to function by binding to promoter regions and opening the chromatin^45^. Expression of tMAC target genes is enhanced through recruitment of Mediator and the tTAFs to those promoter regions^46^. Mutation of any of the tTAFs or tMAC components results in meiotic arrest, where spermatocytes fail to initiate the meiotic divisions^17^.

As the nucleolus was thought to be an unlikely place for Pol II-mediated transcription, it was proposed that the tTAFs indirectly activate spermatocyte transcription through sequestration of Polycomb (and Polycomb Repressive Complex 1) to the nucleolus^17, 19^. However, our finding that active Pol II is localized to the spermatocyte nucleolus/NLB prompted us to reexamine the role of nucleolar-localized tTAFs. We found that the expression of the tTAF Spermatocyte Arrest (Sa) coincides with the recruitment of Pol II to the NLB (Fig 5A, Fig S4A). Although tMAC’s localization to the spermatocyte nucleolus has not been documented, we found that tMAC components Always Early (Aly), Matotopetli (Topi) and Cookie Monster (Comr) also exhibit weak localization to the NLB, while the majority of the signal was seen on autosomal chromatin (Fig 5B, Fig S4B-D).

**Figure 5:**
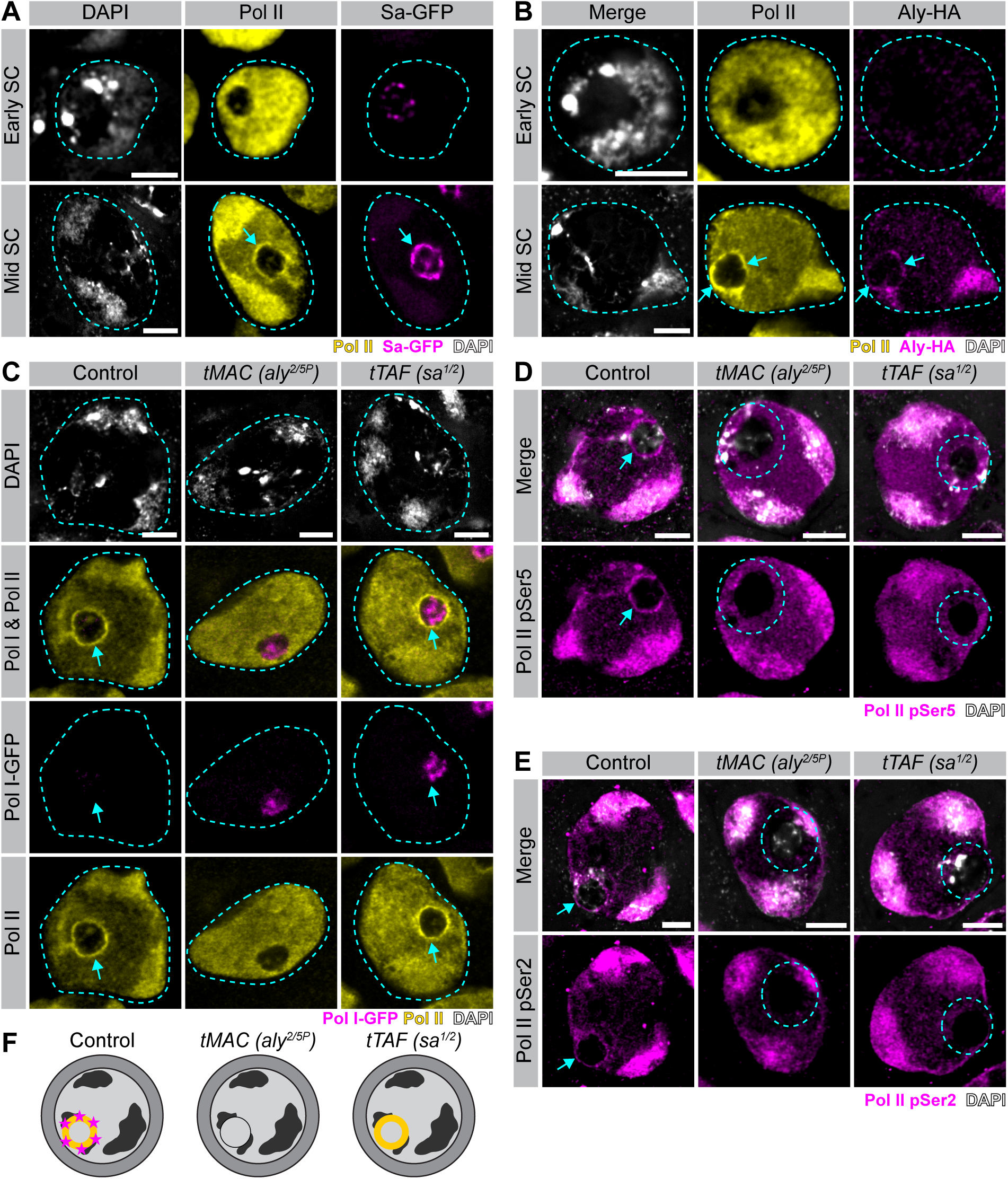
Spermatocyte-specific transcriptional regulators mediate Pol II recruitment to/activation at the NLB (A) Immunofluorescence staining for Pol II (yellow) and DAPI (white) in early and mid spermatocytes expressing Sa-GFP. Spermatocyte nuclei outlined with cyan dashed lines and nucleolar/NLB-localized Pol II and Sa indicated by cyan arrows. Bars: 5µm. (B) Immunofluorescence staining for Pol II (yellow), Aly-HA (magenta), and DAPI (white) in early and mid spermatocytes. Spermatocyte nuclei outlined with cyan dashed lines and nucleolar/NLB-localized Pol II and Aly indicated by cyan arrows. Bars: 5µm. (C) Immunofluorescence staining for Pol II (yellow) and DAPI (white) in mid spermatocytes expressing Polr1B-GFP (magenta) in Control (left), tMAC mutants (*aly^2/5P^*, center) or tTAF mutants (*sa^1/2^*, right). Spermatocyte nuclei outlined with cyan dashed lines and NLB-localized Pol II indicated with cyan arrows. Bars: 5µm. (D) Immunofluorescence staining for Pol II pSer5 (magenta) and DAPI (white) in mid spermatocytes in Control (left), tMAC mutants (*aly^2/5P^*, center) or tTAF mutants (*sa^1/2^*, right). NLB-localized Pol II pSer5 indicated with cyan arrows, NLBs devoid of Pol II pSer5 encircled with cyan dashed lines. Bars: 5µm. (E) Immunofluorescence staining for Pol II pSer2 (magenta) and DAPI (white) in mid spermatocytes in Control (left), tMAC mutants (*aly^2/5P^*, center) or tTAF mutants (*sa^1/2^*, right). NLB-localized Pol II pSer2 indicated with cyan arrows, NLBs devoid of Pol II pSer2 encircled with cyan dashed lines. Bars: 5µm. (F) Cartoon summarizing the state of Pol II at the nucleolus in Control (left), tMAC mutants (*aly^2/5P^*, center) or tTAF mutants (*sa^1/2^*, right). Successful recruitment of Pol II is depicted by the yellow ring and the presence of initiating/elongating phosphorylation marks by the magenta stars.

These results prompted us to test whether the tTAFs and tMAC are required for the recruitment of Pol II to the spermatocyte NLB. Strikingly, the localization of Pol II to the NLB was greatly diminished in mutants for the tMAC component *aly* (*aly*^2^*^/5P^*) (Fig 5C), suggesting that tMAC plays an important role in recruiting Pol II to the NLB. In mutants for the tTAF *sa* (*sa^1/2^*), Pol II was recruited to the NLB in mid spermatocytes (Fig 5C), however, active Pol II markers (phospho-Serine 5 and phospho-Serine 2) were not observed at the NLB (Fig 5D - F), suggesting that the tTAFs are required for the activation of Pol II at the NLB. Consistent with the absence of Pol II in the NLB in tMAC mutants, active Pol II markers were also absent (Fig 5D - F). Coinciding with the defective localization/activation of Pol II at the spermatocyte nucleolus in tTAF and tMAC mutants, we found that Pol I was retained in the nucleolus and rRNA transcription was sustained (Fig S5), implying that the tTAFs and tMAC regulate the conversion from Pol I to Pol II. Together, these results suggest that the tTAFs and tMAC play an important role in the spermatocyte nucleolus transitioning from a canonical nucleolus to an NLB.

### The tTAFs and tMAC are required for proper transcription of the Y-linked fertility genes

Given that Y-linked genes are transcribed at the NLB (Fig 4), where Pol II and the tTAFs/tMAC localize (Fig 3, Fig 5), we examined whether the tTAFs and tMAC are required for the proper expression of the Y-linked fertility genes. Total RNA sequencing of tTAF and tMAC mutant testes showed gross changes in gene expression, as expected from their role as key spermatocyte transcriptional regulators (Fig 6A)^47, 48^. Among these, the Y-linked fertility genes, including those that are likely transcribed at the NLB (Fig 4), are downregulated in tTAF and tMAC mutants (Fig 6A and Fig S6A), suggesting that the tTAFs and tMAC are required for transcription of the Y-linked fertility genes. Consistently, RNA FISH further demonstrated that expression of the Y-linked fertility genes that are likely transcribed at the NLB (Fig 4) (*Ppr-Y*, *kl-2, kl-3,* and *kl-5)* is compromised in tTAF and tMAC mutants (Fig 6B-D, S6B). In tMAC mutants, these genes were either barely detectable by RNA FISH, or had no detectable late exon transcripts (Fig 6D, S6B), indicating failure to complete transcription. In tTAF mutants, transcripts were detectable in the nucleus but functional mRNAs were not detectable in the cytoplasm, apparent by the lack of cytoplasmic RNA granules, which represent mature mRNAs (Fig 6C, D, S6B)^39, 40^. These results demonstrate that the tTAFs and tMAC are required to properly express the Y-linked fertility genes.

**Figure 6:**
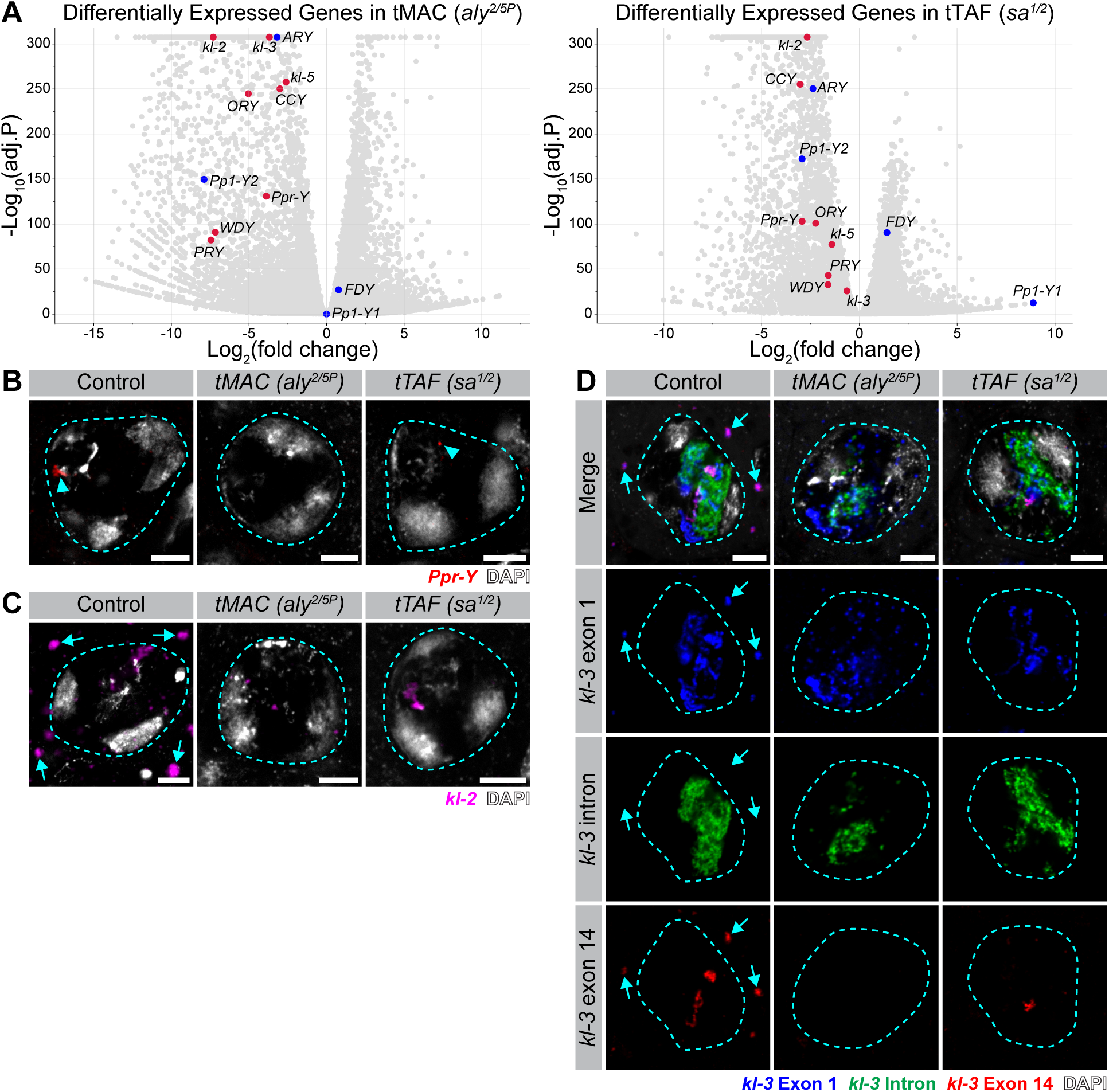
The tTAFs and tMAC are required for proper transcription of the Y-linked fertility genes (A) Volcano plots for tMAC mutants (*aly^2/5P^*, left) or tTAF mutants (*sa^1/2^*, right) highlighting the Y-linked genes. In red are Y-linked genes with gigantic introns while in blue are Y-linked genes with average introns. (B) RNA FISH for *Ppr-Y* in Control (left), tMAC mutant (*aly^2/5P^*, center) or tTAF mutant (*sa^1/2^*, right) mid spermatocytes also stained with DAPI (white). Spermatocyte nuclei outlined with cyan dashed lines. *Ppr-Y* signal indicated with cyan arrowhead. Bars: 5µm. (C) HCR RNA FISH for *kl-2* in Control (left), tMAC mutant (*aly^2/5P^*, center) or tTAF mutant (*sa^1/2^*, right) mid spermatocytes also stained with DAPI (white). Spermatocyte nuclei outlined with cyan dashed lines. *kl-2* mRNA granules indicated with cyan arrows. Bars: 5µm. (D) RNA FISH for *kl-3* (exon 1 in blue, intronic repeats in green, exon 14 in red) in Control (left), tMAC mutant (*aly^2/5P^*, center) or tTAF mutant (*sa^1/2^*, right) mid spermatocytes also stained with DAPI (white). Spermatocyte nuclei outlined with cyan dashed lines. *kl-3* mRNA granules indicated with cyan arrows. Bars: 5µm.

### tMAC is required for the opening of satellite DNA-rich gigantic introns in the Y-linked fertility genes

How is the expression of the Y-linked fertility genes regulated by the tTAFs and tMAC at the NLB? The Y chromosome is largely heterochromatic in most cell types, with the only exception being spermatocytes, where the Y-linked fertility genes need to be robustly transcribed. It has been shown that, when these Y-linked genes are transcribed in spermatocytes, the DNA of these loci loops out into the nucleoplasm, forming lampbrush chromosomes, indicative of robust transcription^49, 50^. These loop structures are called Y-loops^26^. Accordingly, the heterochromatic nature of the satellite DNAs, particularly those in the introns of Y-linked genes, must be overcome to allow for robust transcription. However, the mechanism responsible remains unknown.

DNA FISH against intronic satellite DNAs (AATAT for *kl-3*, AAGAC for *kl-5* and *ORY)*^28^ showed that these intronic satellite DNAs are tightly clustered in earlier germ cells (e.g. spermatogonia/very early spermatocytes) but become dispersed in the developing spermatocyte nucleoplasm (Fig 7A). These organizational changes in satellite DNA likely reflect the opening of chromatin when the Y-linked fertility genes are robustly transcribed as lampbrush chromosomes at the spermatocyte NLB. However, strikingly, these intronic satellite DNAs remain compact and do not spread in the nucleoplasm in tMAC mutants (Fig 7A), suggesting that tMAC is required to open intronic satellite DNA to allow for the transcription of the Y-linked fertility genes. These results imply that tMAC’s proposed function in opening chromatin^45^ may be extended to long stretches of intronic satellite DNA. In contrast, tTAF mutants did not exhibit overt differences in the nuclear organization of intronic satellite DNAs (Fig S7), suggesting that the tTAFs are not required for the opening of satellite DNA-rich introns and instead may function downstream of chromatin opening by tMAC at the spermatocyte NLB.

**Figure 7:**
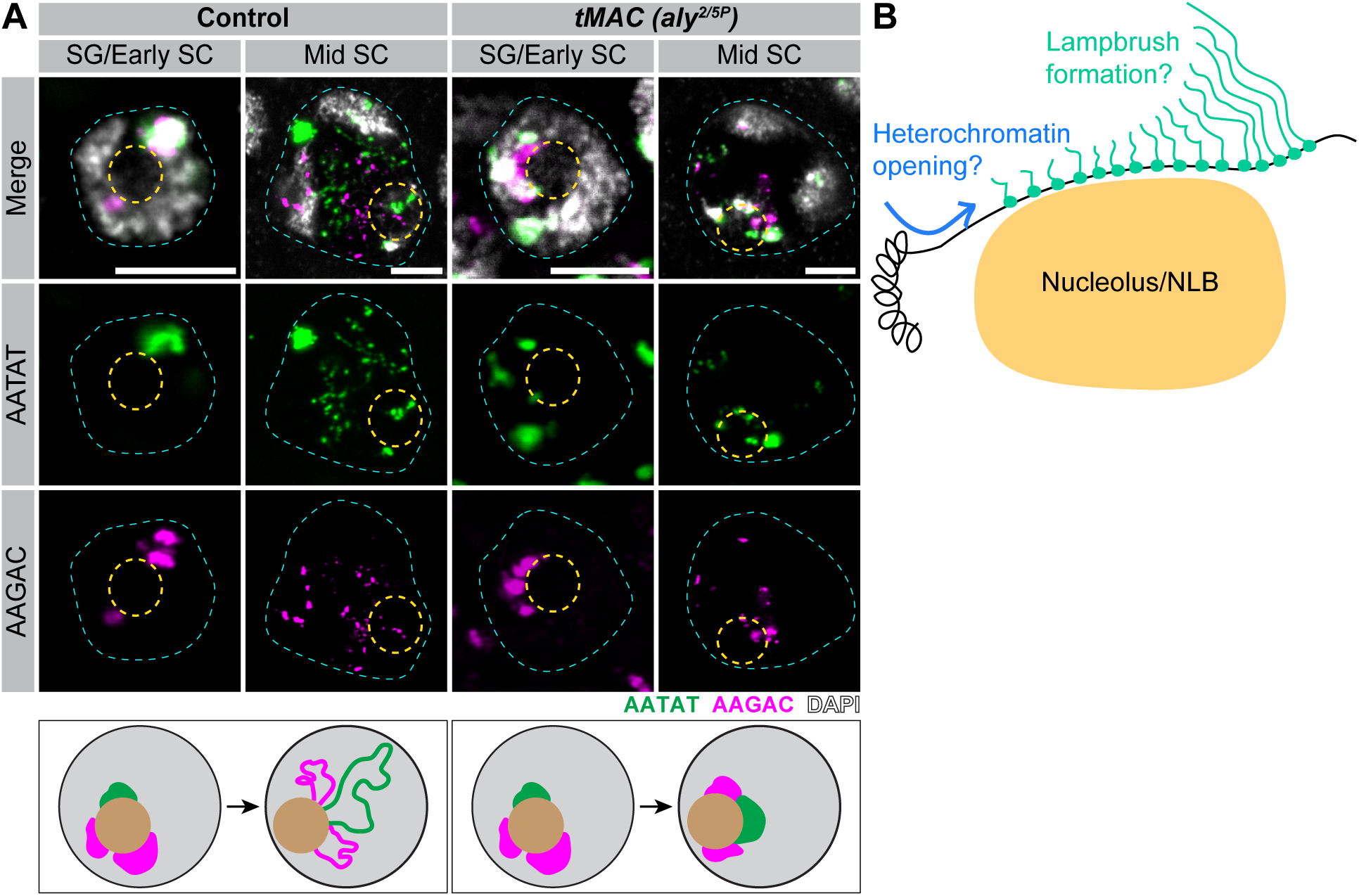
tMAC is required for the opening of satellite DNA-rich gigantic introns in the Y-linked fertility genes (A) DNA FISH for the (AATAT)_n_ (green) and (AAGAC)_n_ (magenta) satellite DNAs in spermatogonia/early spermatocytes and mid spermatocytes in Control (left) or tMAC mutants (*aly^2/5P^*, right) also stained with DAPI (white). The spermatocyte nucleus is outlined in cyan and the nucleolus/NLB in yellow. Diagrams below each image show the degree of compaction for each satellite DNA relative to the nucleolus/NLB (tan). (B) Model for how the nucleolus/NLB may be important for the expression of heterochromatic loci and lampbrush chromosome formation. Heterochromatic loci like the Y-linked fertility genes could be brought to the NLB where NLB-localized transcription machinery (e.g.: the tTAFs, tMAC, Pol II) facilitate the opening of that heterochromatin. These regulators may also promote the robust transcription of these loci and therefore lampbrush formation.

Together, these results suggest that the NLB in *Drosophila* spermatocytes is enriched for transcriptionally active Pol II and is a platform for the transcription of the heterochromatic Y-linked fertility genes (Fig 7B).

## Discussion

The presence of atypical nucleoli, nucleoli negative for rRNA transcription and Pol I, has long been recognized. Termed NLBs or NPBs, their importance in early embryonic development independent of rRNA transcription has been shown^5, 6, 14, 15^, but what role they serve remains unknown. The present study identifies the nucleolus in developing *Drosophila* spermatocytes as an NLB and demonstrates that this NLB serves as a Pol II platform to robustly transcribe heterochromatic Y-linked fertility genes that are silenced in other cell types. Although its universality across species remains to be determined, it is of note that the NLB in mammalian oocytes is known to shutdown Pol I-mediated transcription but contains canonical nucleolar proteins and RNA (but not rRNA)^6, 11^. The presence of Pol II in NLBs has been noted in some cases^31^, but the role of NLB-associated Pol II in transcription has not been suspected. The function of mammalian NLBs remains unclear as does their RNA composition. The present study is the first to demonstrate an active role for an NLB as a Pol II platform. It is tempting to speculate that other NLBs, which were regarded as ‘inactive’ nucleoli or storage bodies, may be active for Pol II-mediated transcription.

Although previous studies proposed that the tTAFs’ function at the nucleolus is to sequester repressive polycomb proteins^17, 19^, our study revises this model, suggesting a more active and direct role for the tTAFs in the nucleolus/NLB. Our results suggest that Pol II is recruited to the nucleolus in a tMAC-dependent manner and activated there in a tTAF-dependent manner. In the absence of the tTAFs or tMAC, the Y-linked genes, which we show to be transcribed at the NLB, were not properly expressed, suggesting that these regulators have a direct role in transcription at the NLB. In particular, tMAC may ‘deheterochromatinize’ the Y-linked genes to facilitate their transcription (Fig 7). In this context, the role of Pc that is recruited to the NLB in a tTAF-dependent manner awaits further investigation. It is tempting to speculate that Pc may be evicted by the tTAFs in the NLB to allow for robust transcription. Together, these results suggest that the tTAFs and tMAC are important for the transition to and activity of the spermatocyte NLB.

This study highlights interesting commonalities in the nucleolar gene expression programs between Pol I-mediated transcription of rDNA and Pol II-mediated transcription of the Y-linked fertility genes. First, both rDNA and Y-linked fertility genes are characterized by their heterochromatic nature. Despite the robust transcription of rDNA, not all rDNA copies are transcribed simultaneously, and many rDNA copies are known to be silenced by heterochromatinization^2, 51^. Additionally, rDNA loci are typically embedded in regions of chromosomes that form perinucleolar heterochromatin^2, 51^. Similarly, Y-linked fertility genes are embedded in heterochromatin, containing introns comprised of heterochromatic satellite DNA repeats and transposable elements^24, 25^. The Y chromosome as a whole is heterochromatic in all cell types except for spermatocytes, and this repressive chromatin state needs to be opened when transcription of the Y-linked fertility genes initiates. Second, both rDNA and the Y-linked fertility genes have many RNA polymerases loaded onto the DNA without an apparent ‘pause-initiation limit’ of polymerase loading, forming lampbrush chromosomes or ‘christmas tree’ structures^49, 50, 52, 53^. Thus, it is tempting to speculate that the nucleolus/NLB may have a unique property that allows for the opening of heterochromatin and the continuous loading of RNA polymerases, overcoming the general constraints of transcription (Fig 7B).

## Methods

### Fly husbandry

All fly stocks were raised on standard Bloomington medium without propionic acid at 25°C, and young flies (1- to 5-day-old adults) were used for all experiments. Flies used for wild-type experiments were the standard lab wild-type strain yw (*y^1^w^1^*). Control flies were either the parental stock or a sibling from the same genetic cross. The following fly stocks were used: *Polr2C-GFP* and *mCherry-Polr2A* (gifts from Patrick O’Farrell)^54^, *sa-GFP*, *p[aly-3xHA-C], sa^1^*, *sa^2^*, *aly^2^*, and *aly^5P^* (gifts from Margaret Fuller)^19, 55, 56^, *Polr1B-GFP* (gift of Eric Wieschaus)^57^, *topi^GFP.FPTB^* (BDSC:56161), and *comr^fTRG0038.sfGFP-TVPTBF (VDRC: v318559)^*.

### Immunofluorescence staining

Testes were dissected in 1X PBS, transferred to 4% formaldehyde in 1X PBS, and fixed for 30 minutes. Testes were then washed in 1X PBST (PBS containing 0.1% Triton X-100) three times, 20 minutes per wash followed by an extended 4 hour long wash in 1X PBST. Testes were blocked for 1 hour in 3% BSA followed by incubation with primary antibodies diluted in 1X PBST with 3% BSA at 4°C overnight. Samples were washed for at least 1 hour in 1X PBST, blocked for 15 minutes with 3% BSA, and incubated with secondary antibody in 1X PBST with 3% BSA at 4°C overnight. Then samples were washed as above, and mounted in VECTASHIELD with DAPI (Vector Labs). Images were acquired using Leica Stellaris8 confocal microscope with a 63X oil immersion objective lens (NA = 1.4) and processed using ImageJ software. For samples stained for Pol II, reagents used were RNase free.

The primary antibodies used were: anti-RNA Pol II (1:100, mouse, Sigma, 05-623), anti-RNA Pol II pSer5 (1:200, rabbit, Abcam, ab5131), anti-RNA Pol II pSer2 (1:200, rabbit, Abcam,ab5095), anti-mod (1:500, guinea pig)^58^, anti-Fib (1:200, mouse, Abcam, ab4566), anti-HA (1:200, rat, Roche, 12158167001), anti-GFP (1:200, chicken, Aves, 1020). Alexa Fluor–conjugated secondary antibodies (Life Technologies) were used at a dilution of 1:200.

### Phase contrast and widefield microscopy

Testes were dissected in 1X PBS and transferred to slides for live observation by phase contrast and widefield (L5 filter cube, Ex: 480/40nm, Em: 527/30nm, Dichromatic Mirror: 505nm) on a Leica DM5000B microscope with a 40X objective (NA = 0.75) and imaged with a Leica DMC4500 CCD camera. Images were adjusted using ImageJ software.

### RNA Fluorescent *in situ* hybridization

#### Single molecule/repetitive transcript RNA FISH

RNA FISH was performed as previously described^59^. All solutions used were RNase free. Testes from 1–5 day old flies were dissected in 1X PBS and fixed in 4% formaldehyde in 1X PBS for 30 minutes. Testes were washed briefly in 1X PBS and permeabilized in 70% ethanol overnight at 4°C. Testes were briefly rinsed with wash buffer (2X saline-sodium citrate (SSC), 10% formamide) and then hybridized overnight at 37°C in hybridization buffer (2X SSC, 10% dextran sulfate (sigma, D8906), 1mg/mL E. coli tRNA (sigma, R8759), 2mM Vanadyl Ribonucleoside complex (NEB S142), 0.5% BSA (Ambion, AM2618), 10% formamide). Following hybridization, samples were washed three times in wash buffer for 20 minutes each at 37°C and mounted in VECTASHIELD with DAPI (Vector Labs). Images were acquired using a Leica Stellaris8 confocal microscope with a 63X oil immersion objective lens (NA = 1.4) and processed using ImageJ software.

Fluorescently labeled probes were added to the hybridization buffer to a final concentration of 50nM (for probes targeting repetitive DNA transcripts) or 100nM (for smFISH probes targeting exons). Probes against the repetitive DNA transcripts were from Integrated DNA Technologies. Single molecule probe sets were designed using the Stellaris® RNA FISH Probe Designer (Biosearch Technologies, Inc.) available online at www.biosearchtech.com/stellarisdesigner. Each set of custom Stellaris® RNA FISH probes was labeled with Quasar 670, Quasar 570 or Fluorescein-C3. Probe information can be found in (Supp. Table 1).

#### HCR RNA FISH

HCR RNA FISH was performed as previously described^41^. All solutions used were RNase free. Testes from 1–5 day old flies were dissected in 1X PBS and fixed in 4% formaldehyde in 1X PBS for 30 minutes and then washed twice, 5 minutes per wash, in 1X PBS 0.1% Tween-20. Permeabilization was achieved by washing overnight in 1X PBS 0.1% Triton X-100 at 4C. The testes were then washed twice, 5 minutes per wash, in 5X SSC 0.1% Tween-20 and blocked with 100ug/mL of salmon sperm DNA for 30 minutes at 37C in Hybridization Buffer (Molecular Instruments). Probes were added to Hybridization Buffer to a final concentration of 10nM. If combining with smFISH or repeat probes, those probes were also added to Hybridization Buffer to their standard concentration (see above). Hybridized overnight at 37C, shaking. Samples were washed four times, 15 minutes per wash, at 37C with Probe Wash Buffer (Molecular Instruments). During the second wash, hairpin solutions for desired amplifiers (Molecular Instruments) were created by individually heating each hairpin at 95C for 90 seconds in a PCR machine and allowing the samples to slowly cool to room temperature. Amplifiers were added to Amplification Buffer (Molecular Instruments) to a final concentration of 60nM. Samples were washed twice, 5 minutes per wash, with 5X SSC 0.1% Tween-20, and then transferred to Amplification Buffer + amplifiers and incubated overnight at room temperature while nutating. Testes were then washed twice, for at least 30 minutes per wash, with 5X SSC 0.1% Tween-20 and mounted in SlowFade Gold antifade mounting media with DAPI (Invitrogen). Images were acquired using a Leica Stellaris8 confocal microscope with a 63X oil immersion objective lens (NA = 1.4) and processed using ImageJ software.

HCR probes designed to span exon-intron junctions were the “V2” HCR probe design while other probe sets were “V3”^60, 61^. Approximately 15bp on either side of the junction was selected for 30bp of target sequence. Probe information can be found in (Supp. Table 1.

### Immunofluorescence staining combined with RNA FISH

To perform IF and RNA FISH sequentially, after fixation, testes were washed twice, 5 minutes per wash, in 1X PBS 0.1% Tween-20, and permeabilized overnight in 1X PBS 0.1% Triton X-100 at 4C. Testes were then blocked for 1 hour in 3% BSA and primary and secondary antibody incubations were carried out as described above. After washing out the secondary antibody, samples were re-fixed in 4% formaldehyde for 10 minutes before being washed twice with 1X PBS 0.1% Tween-20 and twice with 5X SSC 0.1% Tween-20 for 5 minutes each before proceeding to probe hybridization as described above for HCR RNA FISH. For IF/RNA FISH using only single molecule RNA FISH probes, those probes were still added to HCR hybridization buffer but no amplification step was performed. Testes were mounted in VECTASHIELD with DAPI (Vector Labs) or SlowFade Gold antifade mounting media with DAPI (Invitrogen) depending on probes used. Images were acquired using a Leica Stellaris8 confocal microscope with a 63X oil immersion objective lens (NA = 1.4) and processed using ImageJ software.

### DNA fluorescent *in situ* hybridization (DNA FISH)

Testes from 1-5-day-old males were dissected and fixed in 4% formaldehyde + 1mM EDTA for 30 minutes. Samples were washed for at least one hour in 1X PBS 0.1% Triton X-100 + 1mM EDTA and rinsed with 1X PBST without EDTA. Samples were incubated with 2 mg/ml RNase A solution (in 1X PBST) at 37 °C for 10 minutes, followed by washing with 1x PBST + 1 mM EDTA for 5-10 minutes. Samples were then rinsed in 2x SSC + 1 mM EDTA + 0.1% Tween-20 and washed in 2x SSC + 0.1% Tween-20 with increasing formamide concentrations (20%, 40%, and 50%) for 15 minutes each, followed by a final 30 minute wash in 2x SSC + 0.1% Tween-20 + 50% formamide. Hybridization mix (50% formamide, 10% dextran sulfate, 2x SSC, 1 mM EDTA, 1 mM probe) was added to the washed samples. Samples were denatured at 92 °C for 2 minutes and then incubated overnight at 37 °C. After hybridization, samples were washed three times in 2x SSC + 1 mM EDTA + 0.1% Tween-20 for 20 minutes each and mounted in VECTASHIELD with DAPI (Vector Labs). The following satellite DNA probes (from Integrated DNA Technologies) were used: Cy5-(AATAT)_6_, and Cy3-(AAGAC)_6_.

### RNA isolation and sequencing

Total RNA was purified from adult testes (100 pairs/sample) by TRIzol (Invitrogen) extraction according to the manufacturer’s instructions. Libraries were prepared for RNA sequencing using the KAPA Biosystems RNA HyperPrep Kit with RiboErase according to manufacturer’s directions with some modifications. Briefly, 500 ng of total RNA was ribo-depleted by hybridization of complementary DNA oligonucleotides. The set of complementary oligonucleotides was a custom panel designed for Drosophila. This was followed by treatment with RNase H and DNase to remove rRNA duplexed to DNA and original DNA oligonucleotides. The enriched fraction was then fragmented with heat and magnesium, and first-strand cDNA was generated using random primers. Strand specificity was achieved during second-strand cDNA synthesis by replacing dTTP with dUTP, which quenches the second strand during amplification, and the cDNA is then A-Tailed. The final double strand cDNA was then ligated with indexed adapters. Finally, the library was amplified using a DNA Polymerase that cannot incorporate past dUTPs, effectively quenching the second strand during PCR. Libraries were enriched for fragments between 500 – 1000bp with two additional cycles of PCR followed by a size selection using a 1.5% gel on a Pippin Prep (Sage Science) electrophoresis instrument. Final libraries were quantified by qPCR and Fragment Analyzer. Samples were sequenced on a NOVASEQSP, producing 250 × 250bp paired-end reads. Raw sequencing reads in the form of FASTQ files, and the associated gene count matrix are available on the Gene Expression Omnibus (GEO) under accession number GSE332594.

### Bioinformatics analysis of differential gene expression

To analyze differential expression, paired-end reads (250 x 250 bp) were mapped to the D. melanogaster Release 6 (dm6) reference genome using the STAR aligner (v. 2.7.1a)^62^. Counts for fly protein coding genes and lncRNAs (FlyBase Dmel Release 6.38 annotations) were tabulated using featureCounts with the appropriate strand-specific settings^63^. Differential expression of mRNAs was assessed for pairwise contrasts between conditions using estimated fold-changes and the Wald statistic in DESeq2 (v. 1.36.0)^64^.

## Acknowledgements

We thank Drs. Margaret Fuller, Patrick O’Farrell, and Eric Wieschaus, the Bloomington Stock Center, and the Vienna *Drosophila* Resource Center for reagents and members of the Yamashita lab for discussions and comments on the manuscript. We thank Dr. Eliezer Calo for helpful suggestions and Drs. Peter Andersen and Rippei Hayashi for discussions and sharing of unpublished results. We thank the Genome Technology Core and the Bioinformatics and Research Computer Core at the Whitehead Institute for their consultation and aid in designing and performing RNA sequencing experiments.

## Funding

This work was supported by the Howard Hughes Medical Institute (to YMY).

## Author contributions

Conceptualization: JMF, JIP, YMY

Data curation: RL

Formal analysis: JMF, RL, YMY

Funding acquisition: YMY

Investigation: JMF, JIP, RYL, AA, YMY

Supervision: YMY

Validation: JMF Visualization: JMF

Writing -- original draft: JMF, YMY

Writing – editing: JMF, YMY

## Data Availability

All relevant data are within the paper and its Supporting Information files as well as at the GEO under accession number GSE332594.

## Competing interests

The authors declare no competing interests.

## Supplementary Figures

**Figure S1:**
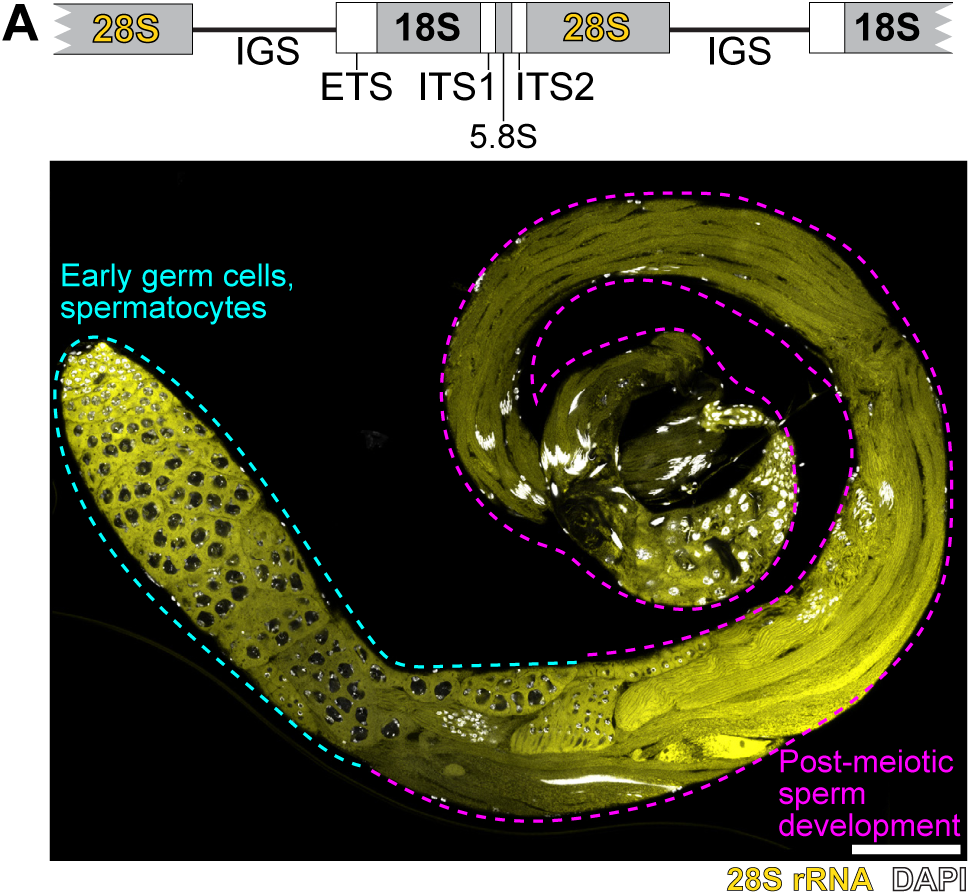
rRNAs are abundant in post-meiotic sperm development (A) Top: Diagram of rDNA akin to that in Figure 3A. Bottom: RNA FISH for 28S rRNA (yellow) in a whole testis also stained with DAPI (white). Portions of the testis containing early germ cells and spermatocytes are outlined with cyan dashed lines while post-meiotic stages are outlined with magenta dashed lines. Bar: 100µm.

**Figure S2:**
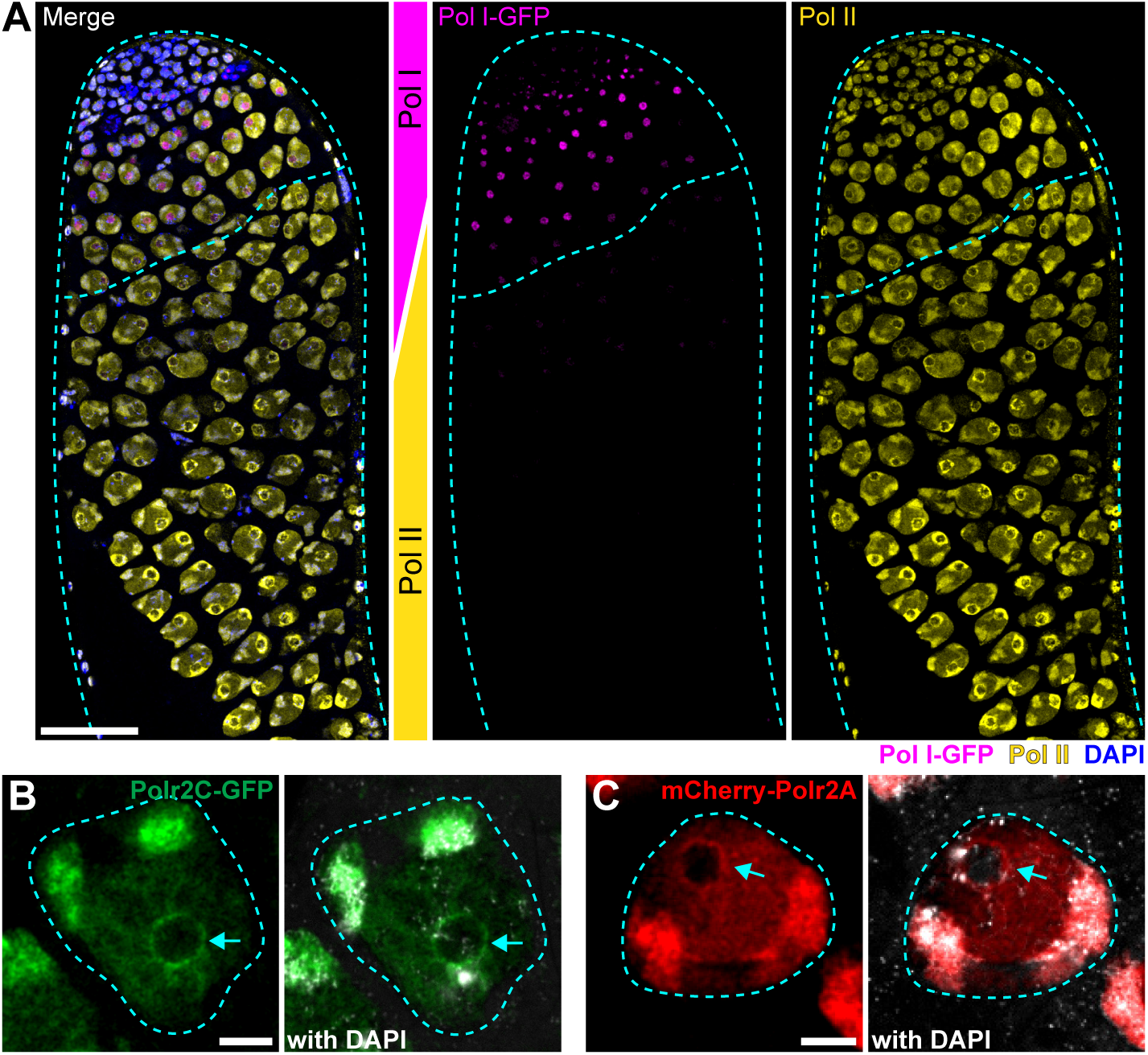
Pol I disappears from the nucleolus concomitant with Pol II recruitment (A) Immunofluorescence staining of the apical tip of a testis through the end of spermatocyte development (cyan dashed outline) showing Polr1B-GFP (magenta) Pol II (yellow) and DAPI (blue). Horizontal cyan dashed line indicates the transition point where Pol I and Pol II have switched occupancy of the nucleolus/NLB. Bar: 50µm. (B) Polr2C-GFP (green) expression in a mid spermatocyte nucleus (cyan dashed outline) with DAPI (white). NLB marked by cyan arrow. Bar: 5µm. (C) mCherry-Polr2A (red) expression in a mid spermatocyte nucleus (cyan dashed outline) with DAPI (white). NLB marked by cyan arrow. Bar: 5µm.

**Figure S3:**
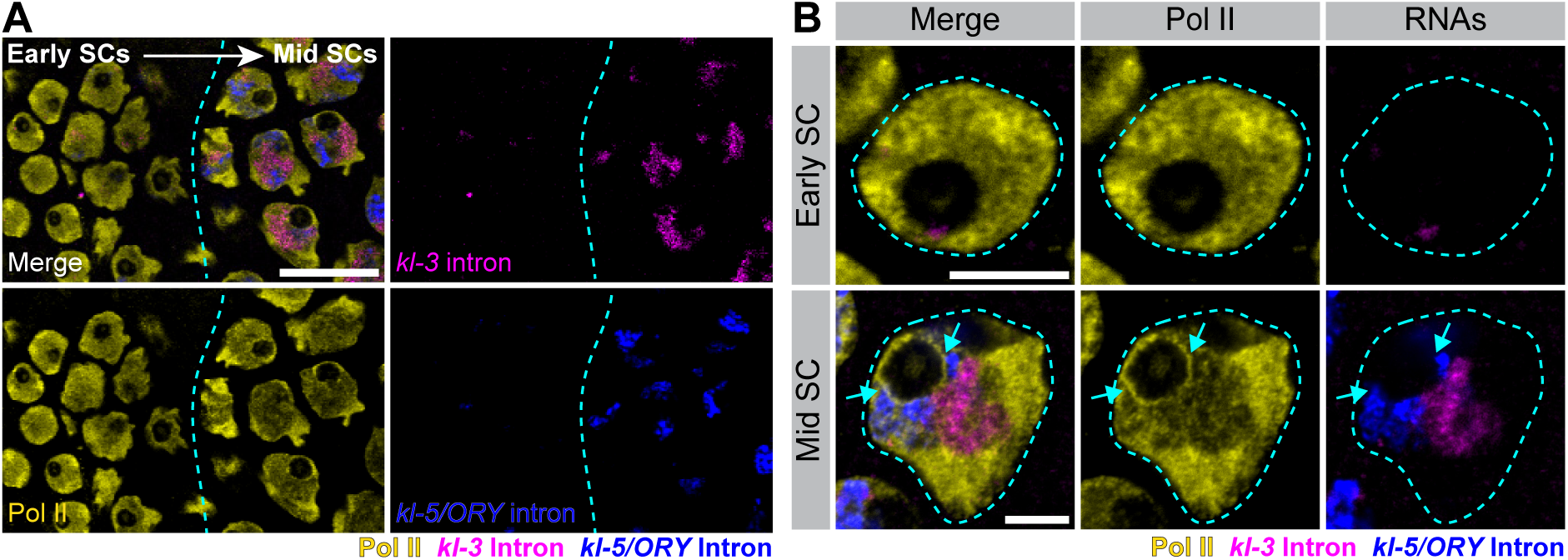
Localization of Pol II to the NLB coincides with the burst of Y-linked gene expression (A) Immunofluorescence staining for Pol II (yellow) with RNA FISH for the intronic repeats of the Y-linked genes *kl-3* (magenta) and *kl-5* and *ORY* (blue) in early-to-mid spermatocytes. Vertical cyan dashed line indicates the point at which Pol II becomes nucleolar. Bar: 20µm. (B) Immunofluorescence staining for Pol II (yellow) with RNA FISH for the intronic repeats of the Y-linked genes *kl-3* (magenta) and *kl-5* and *ORY* (blue) in early and mid spermatocytes. Spermatocyte nuclei are outlined with cyan dashed lines. Cyan arrows mark nucleolar/NLB-localized Pol II. Bars: 5µm.

**Figure S4:**
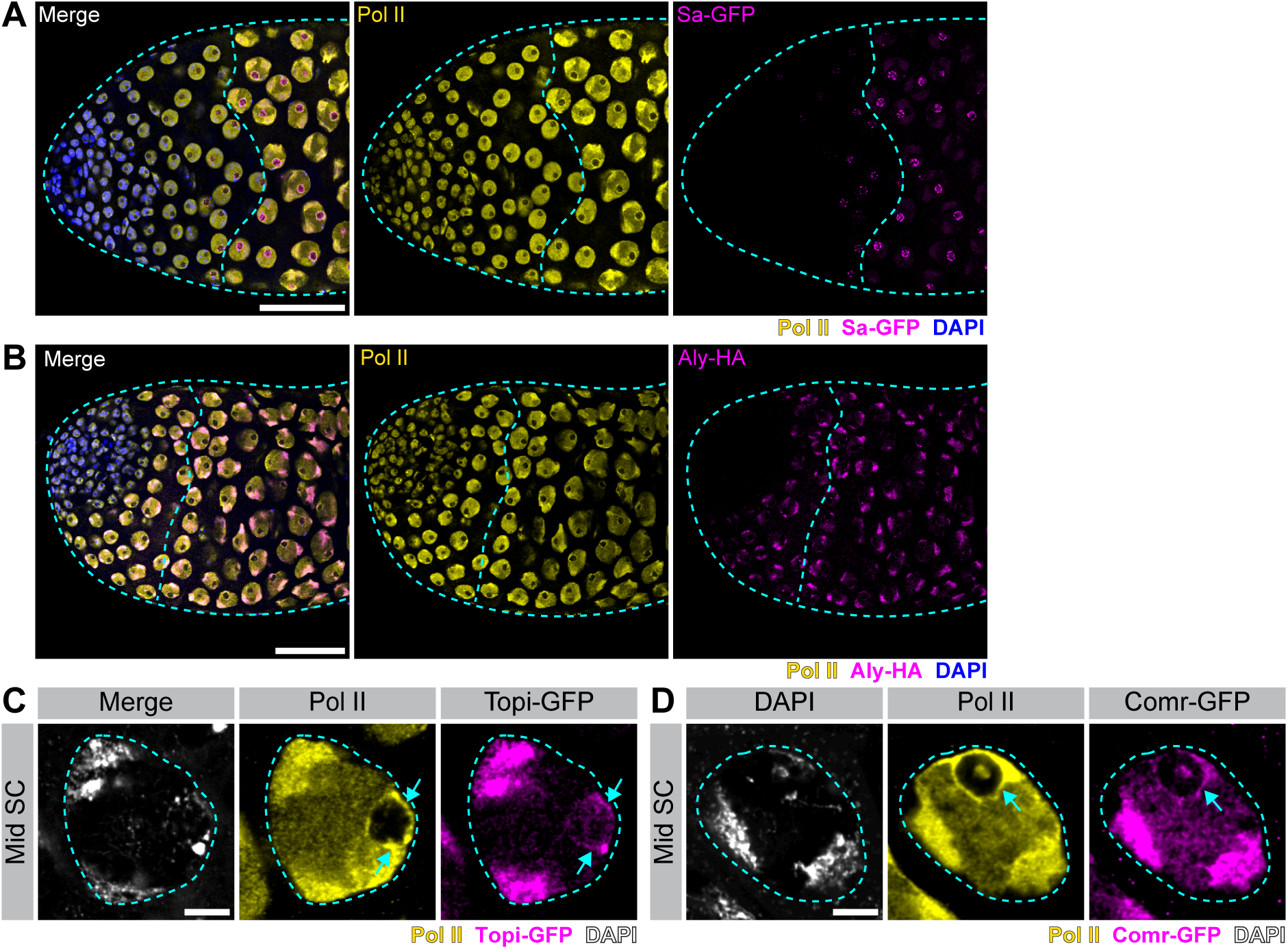
The tTAFs and tMAC localize to the spermatocyte NLB (A) Immunofluorescence staining for Pol II (yellow) and DAPI (blue) in the apical tip of a testis (cyan dashed outline) expressing Sa-GFP (magenta). Shown are germline stem cells through mid spermatocytes. The vertical cyan dashed line indicates the point at which Pol II becomes nucleolar/NLB-localized. Bar: 50µm. (B) Immunofluorescence staining for Pol II (yellow), Aly-HA (magenta) and DAPI (blue) in the apical tip of a testis (cyan dashed outline). Shown are germline stem cells through mid spermatocytes. The vertical cyan dashed line indicates the point at which Pol II becomes nucleolar/NLB-localized. Bar: 50µm. (C) Immunofluorescence staining of Pol II (yellow) and DAPI (white) in a mid spermatocyte expressing Topi-GFP (magenta). Spermatocyte nucleus outlined with a cyan dashed line. NLB-localized Pol II, Topi indicated by cyan arrows. Bar: 5µm. (D) Immunofluorescence staining of Pol II (yellow) and DAPI (white) in a mid spermatocyte expressing Comr-GFP (magenta). Spermatocyte nucleus outlined with a cyan dashed line. NLB-localized Pol II, Comr indicated by cyan arrows. Bar: 5µm.

**Figure S5:**
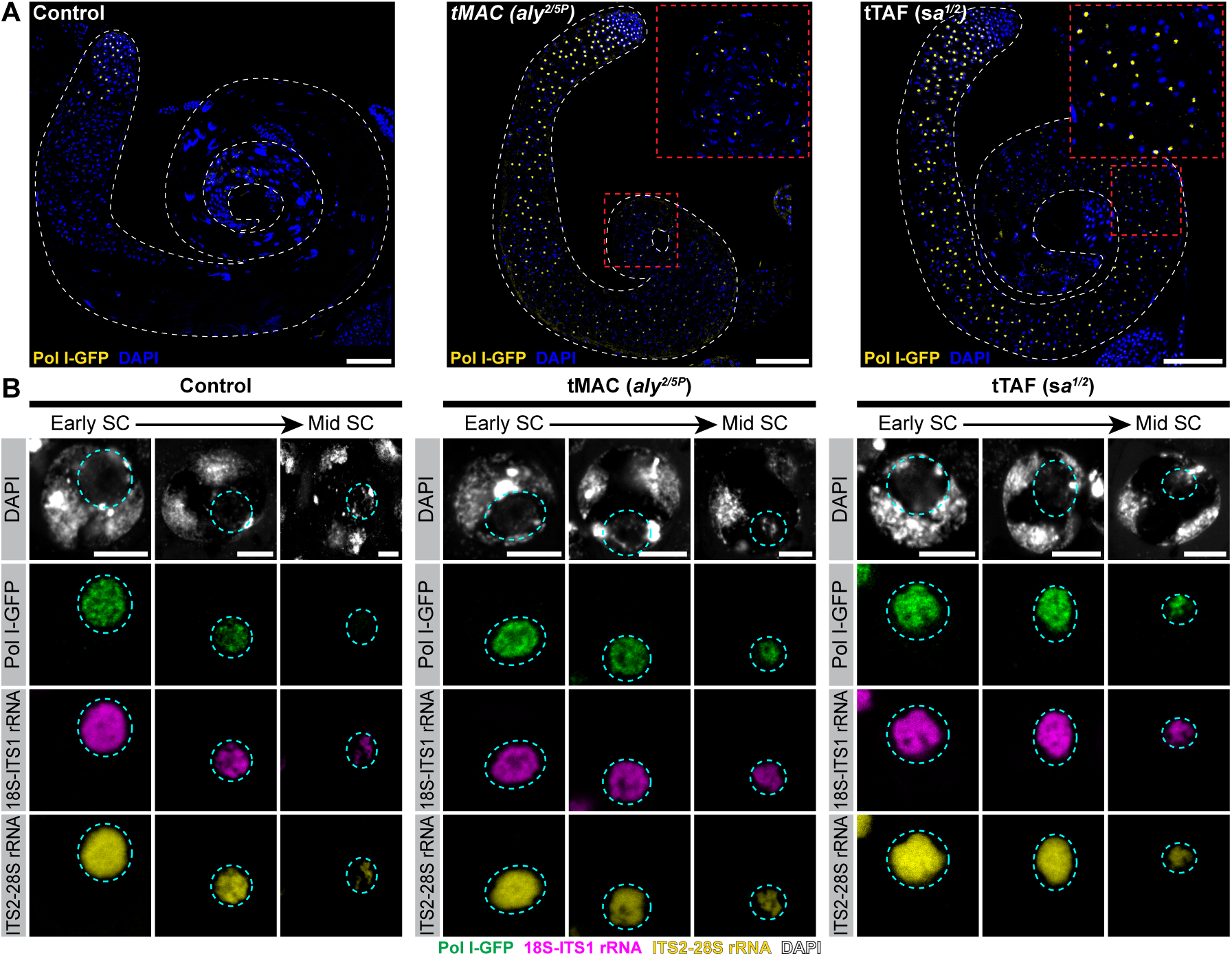
Pol I and rRNA expression are retained in the nucleolus of tMAC and tTAF mutants (A) Expression of Polr1B-GFP (yellow) in a whole testis (white outlines) in Control (left), tMAC mutants (*aly^2/5P^*, center) or tTAF mutants (*sa^1/2^*, right) also stained with DAPI (blue). Insets show arrested spermatocytes in tMAC and tTAF mutants expressing Polr1B-GFP. Bars: 100µm. (B) RNA FISH for unprocessed 18S-ITS (magenta) and ITS-28S (yellow) rRNAs over spermatocyte development in spermatocytes also expressing Polr1B-GFP (green) and stained with DAPI (white) in Control (left), tMAC mutants (*aly^2/5P^*, center) or tTAF mutants (*sa^1/2^*, right). Nucleolus encircled with cyan dashed line. Bars: 5µm.

**Figure S6:**
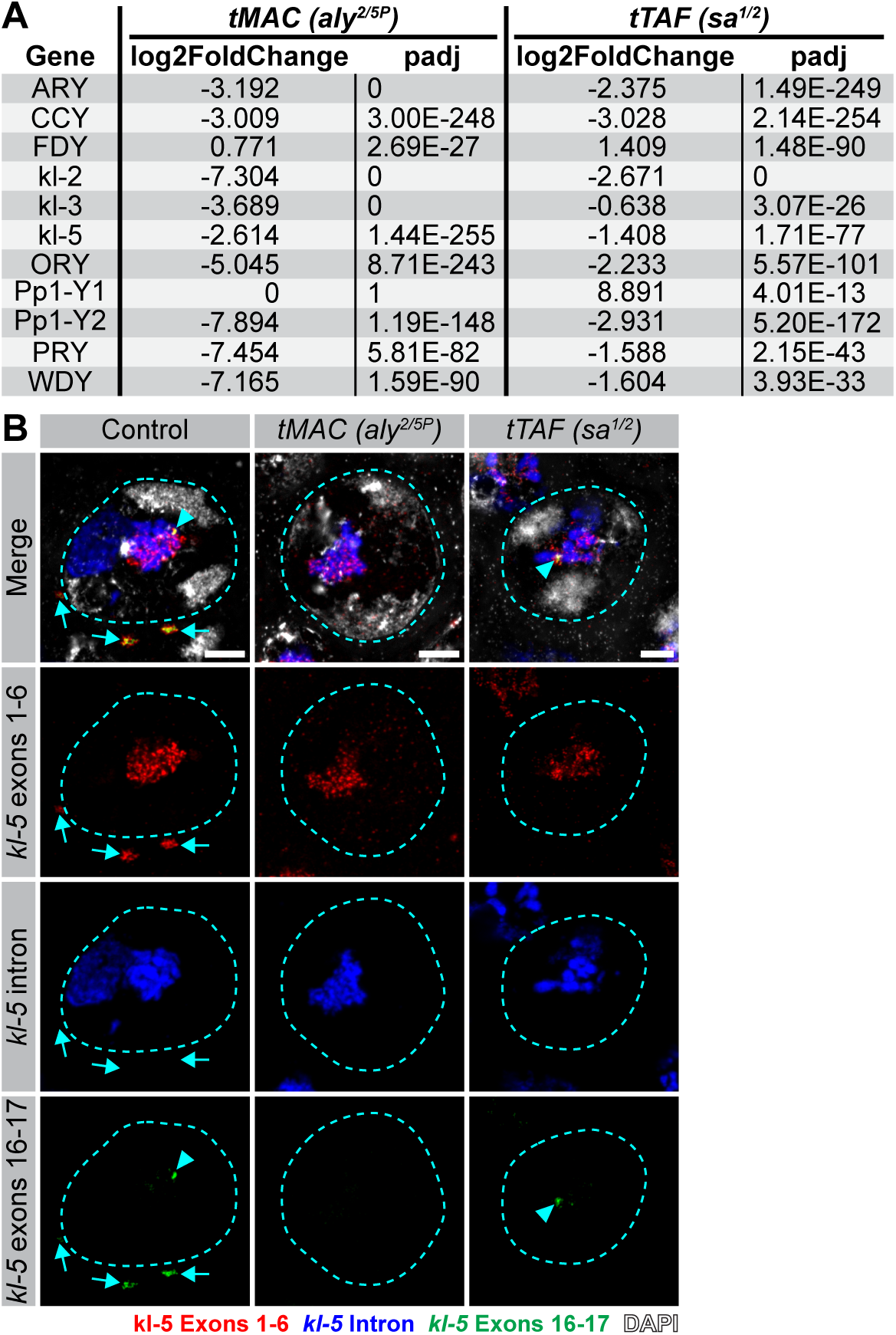
The tTAFs and tMAC are required for proper Y-linked gene expression (A) Table with log2FoldChange and p-adj values corresponding to the Y-linked genes highlighted in the volcano plots in Figure 6A for tMAC mutants (*aly^2/5P^*, left) or tTAF mutants (*sa^1/2^*, right). (B) RNA FISH for *kl-5* (exons 1-6 in red, intronic repeat in blue, exons 16-17 in green) in Control (left), tMAC mutant (*aly^2/5P^*, center) or tTAF mutant (*sa^1/2^*, right) mid spermatocytes also stained with DAPI (white). Exons 16-17 nuclear signal highlighted with a cyan arrowhead. Spermatocyte nuclei outlined with cyan dashed lines. *kl-5* mRNA granules indicated with cyan arrows. Bars: 5µm.

**Figure S7:**
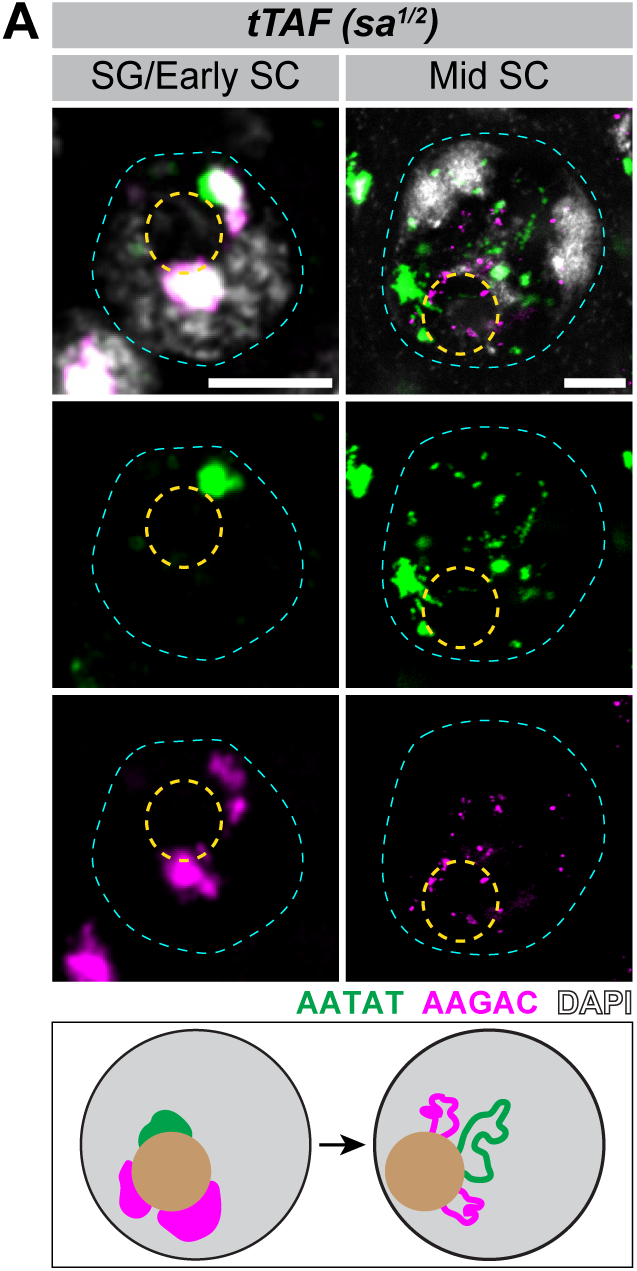
Y-linked satellite DNAs are opened in tTAF mutants (A) DNA FISH for the (AATAT)_n_ (green) and (AAGAC)_n_ (magenta) satellite DNAs in spermatogonia/early spermatocytes and mid spermatocytes in tTAF mutants (*sa^1/2^*) also stained with DAPI (white). The spermatocyte nucleus is outlined in cyan and the nucleolus/NLB in yellow. Diagrams below each image show the degree of compaction for each satellite DNA relative to the nucleolus/NLB (tan).

## Notes

### Competing Interest Statement

The authors have declared no competing interest.

