## supplementary file for "Developmental conversion of the nucleolus into an RNA Polymerase II transcriptional platform in *Drosophila* spermatocytes"

Supplementary Table 1: RNA FISH probes

Single molecule RNA FISH probe sets

| **Probe Target** | **Fluorophore** | **5’-Sequence-3’** |
| --- | --- | --- |
| *kl-3*, Exon 1 | Quasar® 670 | taacattcctttctggatcc, cgcgaaacgccaaagagttt, gcagcacgctttaacatgtt, ttggtcacttacactaggtc, tatcgtcttctttgttggtc, cctcatttctcgaagtaact, caggcttgaaatccgttgtt, aaacataccgctggtttggg, atacccaacaaatctgcaca, gttactatttcctcaggatc, ttgctttcatccacaatacc, accatttacattctcaacat, gggacctttttcctcaaata, cgttacttatcattatggcc, cggaatcagttggatatcct, tttaagcttttcctggtagc, agagcgttgaatttcagtgt, actagatccgacctcaaaca, gtcgatatacaacagtccac, accgaacggttatcaatcga, aaatattgccacctcatcac, aagcaagagtttcgctcttc, cggctttaaaacgtgatcca, caaattcggtcacagcttct, tgagtttgctctttttctgc, acagttgcttcagatgattt, cctttaaatagttcatgcga, tttgccatctcacaagtaat, gtatttttgaactggcttca, caaccagtcgaactcgtgta, gccattgctcaaaataacgt, gtataccttgaatttgtcga, gtatttgttttccctctact, ctacatcgggtgtatctctt, ccaattgaccaacatttgca, tatatctggccaacatacgc, gtaacaaactccgttatggt, gtggttattaaatgctcggg, agataatgtcaagcaatcct |
| *kl-3*, Exon 14 | Quasar® 570 | gtactttgacatagccatgg, aagatttgcctttaagggca, tgatatttagcctcttgcac, ctgcttcttgtagatcactt, cgttttctttttgttgcagt, tttttggcctcgtctaatac, aaccaccaataagagcggtt, ttcagtccatcggatttttt, cggtcggtctcacttttaaa, ggagaataacatctccgacc, tcttgattaaatggtcccgt, atgttccactcaccaatttg, ttgacagttcatctgttggt, ttttaatccacacttttccc, taatcggaattcccatgcta, taactcctctgcaacatctt, aaggttgtccaagcaaggat, tccaatccacttctttatca, gattgggtagctttgttgta, cgcgcaaatatttcaggtgt, ccacgcattgtcacagtaaa, gaacccgttcatcttctaat, tttcatgtttccagtcacag, ttcctttcgtagtggacaac, cctcaatcactgtaacgtcg, tccttaacttcaatggcagt, cagcgtttatttttgcttct, acactacctcttgtagcaac, gcatcaaagcgctcaaggaa, tcctctagatttgtagcgat, ttaactcaagctggcgatca, cgcaccgccttttataaata, caccgaaacggaacaggagg, tataccacgtaaaccaagcc, ccatacaccacgaacgaaca, agtttagtactactggctcg, aatcattggcataagttccc, agctcattttttttggcgag, cttggcccatagaaatagga, cgtcttctaggcaggataat, tcctaagtgacagttttgca, actgtaagctctaccatgta, agcaggtggttcattagtat, tttttagtccagcacgtatt, tggagattgcgaatagtcca, tgcataccagtcggatgaat, tcctgctccttaaaactgta, cgtttgccatgtcatcaata |
| *kl-5*, Exons 1-6 | Quasar® 570 | cttcttttccttttcgtcag, aaaaactccggacggttgtc, ttgtcttggttaggtagttc, ccacttatccagcttaagac, cctaaactcgttggttgtta, atacgtttcttgttgggatt, ccaccggaattgattgtgaa, ctggaaagctgtaggatgga, agcaactttataccgtggtc, gaagtggtgttaagtaccga, atttgctaatgggttcggta, atcccacttgatttactgtg, tctgtcttcatatcgtttgc, tccatttcgcatttcttgag, aacgagacctttcatctggg, catgacttcgtccatgcaaa, gcttcataagagggttaacc, caagccactttacaaccata, tttacgaggtcttcaacgga, tgcctctggtagtggaaaat, caagttctcaaggttttcca, agtctttatacgcctatctc, tctgcgataagaatgctcga, acttagttatgtctctagcc, gtccatatcgtttgtttcaa, gtcgtatatctgatcggttc, cactaatccaattgtcagca, tgaaaataccgcgagtgacc, gttctgttgacacagtcgat, ggaatatggagtctggttct, ttataggcttcgtcgacatc, tgtaatactctaggtgctgc, cagtcctgcgagttaaatgt, atcgtccgaatatttcatcc, tcttttagctcttccaatcg, ccccataacaattttttcca, aagggcgaaagttatctgcc, tgctgttgtactcttcaaga, aatatttgtccactcacggt, acagagaattttctgcgtcc, tggccttgaagcttaaacga, tgaaacgataccatgcgctc, atgtgtgtcttcaagacctt, gcgaagaagtaaggagcctg, gggttcgacatgttttttga, atcatccacaatgacatgca, taacgcacatgatctcttcc, cgagcgccgtaaaaacgttt |
| *kl-5*, Exons 16-17 | Fluorescein | ttgtggtccgaatttcctac, taaacgggtagctacggttc, tgatattgtcaaatcgccca, ccaggtaattgtacagcaca, caaggtactctggtattggc, ccatacataatctctccgaa, ccagtcgtctgtaatgtgac, gataagttcggcataagcgg, agctctggctgcataaattc, aaaccctggcaatactctag, atttaagtattcctggagcc, atgtaattgtggtagccagt, gagatggactttcagacgga, gcatttgaatgaaggccgta, cgtagttaggaacccaatct, attcggaagagtcgttcaga, tctcggctgcaactcgaaaa, aaactgtctcaccaccacta, gttttttataatgtcttcct, aatggtgtgggagttttgtc, cgcgacccattagttctaaa, atgtatggacttcggtcttc, cgctcacactcttgaaatgc, gcttcaattcagtcataaga, aggtcaagctcattcagaga, agtcaattctcctttaaggc, ccattaaatcctccataact, gcacttggtccatgtaaaga, aaccaggattgtagccctag, cagccgcaacattaaatcgg, aggcatgcgaaagtcggcaa, atcctgccaaccaaattgat, aggagcgactgaggattaaa, cgtctgttgcatgatagctg, gacacattctatcgagaggc, ttccactttttggtgacatc, ttgatataggcaccctctcg, tgctccctccatgaaaagac, aatggttcccattttcatgt, gttcctttaaaaaggcgtct, agaacaggcatagcaggaaa, cttgggttactgcctttatg, gacatttttaatatcctgct, cggattttgtaaactgggca, caaacgaaggtgggtcctcg, ctctcggcttttcaagttaa, ctagagtccacttacttgct, tgcaagagaagacaaacccc |
| *Ppr-Y* | Quasar® 570 | atttatttgatcgtcttccat, aataacttcatccacaagggg, tattccaggttcaatatctgg, agctttcaatcaaggaacggt, caccacgatgaacgtgtttta, cagttgatgtaaacgtcttcc, atttgttccaatacaacaggc, ttaaactccaaacgcattgtt, atccataagtgatcaatacgt, taaggcaaagcttggtcagat, aatggtctctattttgttgca, ggtgtcaagattttcgatctt, gacagaacttccaaatttact, ctcaatagcttcaattttgtt, agcatggttaacatatcgata, tatttccaaggctgataatta, tccttcaactgtgtcaattaa, ttcataaaccgaaatcgttca, gataggattaccttccaaatt, ctgatatgtactttaacaggc, cggagttcactcttaatgaaa, agtataaattacaggcttctg, tcttcaatttcccgaatctcc, cttgtatctccttttcttgtt, tctgattgttcacgttcaaga, aacttgaggctaatcgttttg, tgatgaccatccaaatgttca, cacgccaaagtgagtcaaaca, caacaagcatcagcacacgac, atatccttatcgtattcgtcg, gtaaatttcttgcgtaagttc, gtaaatctttctaagccaagt, tctcgaatttcttcatcgcgc, tatttgaagttcttcttggcc, acataagctccgcaatttttc, atgcttgagattgttctacaa, tcgagtcaatatttcggtcat, gtcttgaagtgcaaatttcgt, tcggagttcaataggaacgtc, ttaaaatggcgtcacggtcat, ctcgctcatctaatcgcaaag, gtctacaaattcctttgttcg, ctgtttagttggtctatcata, aatttttgaccgatgtcgttc, catcggtcatcatttctacaa, cttacatcgtgtggtaaggat, acataatcttcggctatcagc |
| *18s* | Quasar® 570 | tataactactggcaggatcaac,catggcttaatctttgagacaa,  tcacttttaattcgtgtgtact,actgatataatgagccttttgc,  ctgttaacgatctaaggaacca,agaattaccacagttatccaag,  aggttcatgttttaattgcatg,tagcctaataaaagcacacgtc,  aatataacgatcttgcgatcgc,atacgatctgcatgttatctag,  acatttgaaagatctgtcgtcg,gtcctagatactaccatcaaaa,  gatatgagtcctgtattgttat,agtgtactcattccaattacag,  caattggtccttgttaaaggat,ccgcaacaactttaatatacgc,  agcacaagttcaactacgaacg,acaattgtaagttgtactaccc,  atataagaactccaccggtaat,tgcaggtttttaaataggagga,  cccacaataacactcgtttaag,tgctttaagcactctaatttgt,  cacagaatattcaggcatttga,cagaacagaggtcttatttcat,  cctcttgatctgaaaaccaatg,ccaaactgcttctattaatcat,  ttaagttagtcttacgacggtc,aacatctttggcaaatgctttc,  ctctaactttcgttcttgatta,tcgtttatggttagaactaggg,  gagagagccataaaagtagcta,aattcctttaagtttcagcttt,  aatctgtcttacacacttatgt,ccatagattcgagaaagagcta,  atcactccacgaactaagaacg,ttcgttatcggaattaaccaga,  caccataatcctgaagatatct,gaatgaaggctacataagcttc,  acacaataagcattttactgcc,gctccacttacataaacacatt,  gtgtccttataatgggacaaac,gcaatttgtccatttaagaagc,  ctgttattgctcaatctcatta,ggtctaggaaatacacgttgat,  ttcacaatcccaagcatgaaag,gaattccaagttcatcgtgaac,  caatgcgagttaatgactcaca,taattcaatcggtagtagcgac |
| *28s* | Quasar® 670 | aatatgcttaaattcagggggt,agacaaagtgacttagtgctga,  tagaatattgacgctccataca,cacgactctttggaaacatcat,  tagttttagatggagtttacca,cgtactgtttaactctctattc,  ttttccataacggatattcagg,tgacttcaacctgatcaagtat,  tgagggaaacttcggaaggaac,taatcattcgctttaccagata,  gttcttacccatttaaagtttg,cactgggcacattatatcataa,  gtaacattacgtttggttcatc,agcggtaatgtataggtataca,  ctcattcgattatttgctacta,ttccctttcgttcaaaattatt,  cgaggcccatatttaataacaa,agaaaactcttccgatatctct,  gacacaacagtatatgtcatgc,ttttcaaggtccgaggagaaaa,  acctacattattctatcgacta,ccgaagttacggatctaatttg,  atgttattgtttcccaatcaag,gaaactctatttacccagaacg,  actaaactatccggggaacaag,ggtttaacccaaatagtattct,  tttaattagacagtcggattcc,gaaatcacattgtgtcaacacc,  ttcttcactttgacattcagag,ctaccttaagagagtcatagtt,  cactaattagatgacgaggcat,aagtaggaatctcgttaatcca,  ttagagtcaagctcaaaagggt,ataccaaaccgaggtctaatat,  ggaaatgatacacgttccattt,tactttgatcaagaagcttgca,  tgaattggatcatacctgagta,tccgttattatttgagagatgt,  tgtactgaacaccgagatcaag,ttttctgacacctcttgttaaa,  atcaaaaagcgacgtcgctatg,ggtgaattttgcttcacaatga,  gtaaaactaacctgtctcacga,cgcaacaacgttttgtcattag,  gtttagaagccatacaatgcaa,ctcccaatttactatgttacaa,  ctaggctttgtcattgtattaa,cctgttacaaaagtcgtttaca |

**FISH probes for repetitive sequences**

| **Probe Target** | **5’-Sequence-3’** |
| --- | --- |
| (AATAT)n | Alexa488-ATATTATATTATATTATATTATATTATATT  Cy5-ATATTATATTATATTATATTATATTATATT |
| (AAGAC)n | Cy3-AAGACAAGACAAGACAAGACAAGACAAGAC  Cy5-AAGACAAGACAAGACAAGACAAGACAAGAC |
| 18S-ITS1 | Cy3- AACGGTAAGGATATTATACAATAATGATCC-Cy3 |
| ITS2-28S | GAGTTGAGGTTGTATATAACTTTATCTTGC-Cy5 |

**HCR RNA FISH probes**

| **Gene** | **Target** | **Sequence** | **Amplifier** |
| --- | --- | --- | --- |
| *kl-3* | intron1 - exon2 | CCTCGTAAATCCTCATCAATCATCCAGTAAACCGCCatataGATT  TCAGCCCAGCGctagaaacaaaaaaa | B2 |
| *kl-3* | intron3 - exon 4 | CCTCGTAAATCCTCATCAATCATCCAGTAAACCGCCatataCCACT  TCTTTTATTTTctaatttggaaaaat | B2 |
| *kl-2* | CDS | GTCCCTGCCTCTATATCTttAGTACCTCGCTTTATCCAGTAGTCA  TTAGTTTGCTAAACTTAATAAAAGTttCCACTCAACTTTAACCCG  GTCCCTGCCTCTATATCTttCAATGAATTATATGCTTTTAACCAT  GTCTCTAGCCCATGCCGCTAATGGTttCCACTCAACTTTAACCCG  GTCCCTGCCTCTATATCTttGAAACATTTGTATTTGGTCTAAATA  CAAAAGCATCAGGCTTATCAAAATTttCCACTCAACTTTAACCCG  GTCCCTGCCTCTATATCTttATATTAGCCTCATTTATGTTGTTAT  TACTTGCTGGGTTCACTACAATTTTttCCACTCAACTTTAACCCG  GTCCCTGCCTCTATATCTttGGATCGACACCTGGTGATAATACGA  TCTGATAGTGATATGAGAGATTGGGttCCACTCAACTTTAACCCG  GTCCCTGCCTCTATATCTttTTGGGCGCTTGTCCCAGTTTATCTA  TGCTCACTTATCCACCATGGTGCCGttCCACTCAACTTTAACCCG  GTCCCTGCCTCTATATCTttCTTGTCGGGCCGCATCAATTTCTAC  ATGCGCGTTCCGATGCTGGTTTGTAttCCACTCAACTTTAACCCG  GTCCCTGCCTCTATATCTttATGGTAAAGCTCTGCCGCAAAATTG  AATTTTAATAACCTTTCTCCACTTTttCCACTCAACTTTAACCCG  GTCCCTGCCTCTATATCTttTTTAATTAATAAAGACCATTTTACA  CTCCAAAGTTGCTGGTATTAAAAGAttCCACTCAACTTTAACCCG  GTCCCTGCCTCTATATCTttTAAACTTTTCCGTAATTTTCTATGG  TTTTCCTGCTTTGGAGCGACTATTCttCCACTCAACTTTAACCCG  GTCCCTGCCTCTATATCTttCTAATTCAAGACAAATTATTTCATC  CCTCCAAGTCCGCACGAGCTGTAGCttCCACTCAACTTTAACCCG  GTCCCTGCCTCTATATCTttAGAATGGCTTCTGTGCTTATAACCA  GAAAATACATAGGCAATATCGCGTTttCCACTCAACTTTAACCCG  GTCCCTGCCTCTATATCTttTTGTTTGTTATTTCGAGAAATGAGA  AAGTTTATTTCTCCAGTACTTAGCAttCCACTCAACTTTAACCCG  GTCCCTGCCTCTATATCTttGACTGTGAAAAGTAACTTCAAAATG  AAAATGGTGGAACCTTTGAAGGACAttCCACTCAACTTTAACCCG  GTCCCTGCCTCTATATCTttGAGTTCGACTGGAAATTGTTTGTCT  TTAAGTTTAATAAAACAAACCGACTttCCACTCAACTTTAACCCG  GTCCCTGCCTCTATATCTttGTGTCTCCAGTAGGAACAATAATCT  TTTGAAACAACATATTCATAGCGAAttCCACTCAACTTTAACCCG  GTCCCTGCCTCTATATCTttAACTCCATTTAACTCATTTGTCCTT  TATTTCTAGCAATTTACAAAGCGACttCCACTCAACTTTAACCCG  GTCCCTGCCTCTATATCTttTTATTCGCATAATAGAACTTAAAAC  TCTGAGTCGGCTCTTCATCTCCACAttCCACTCAACTTTAACCCG  GTCCCTGCCTCTATATCTttCTCGCAACATTCATATCTTTCATTG  AAAAGGGGTAAATCATTAGCTGTTAttCCACTCAACTTTAACCCG  GTCCCTGCCTCTATATCTttGCTGCCATTATAGACATTATTTGTT  ATAAGCTCCAACGCCTTTGTAGAAAttCCACTCAACTTTAACCCG  GTCCCTGCCTCTATATCTttCTGCCTTATTACACATAGTTCCGAT  ATAGCCATACCAGTGCTCTGTGTTTttCCACTCAACTTTAACCCG  GTCCCTGCCTCTATATCTttGCATCTCTTTCATAACAACAAGCAT  TTTTAAGAGATCCGCGAGTAAGTTTttCCACTCAACTTTAACCCG  GTCCCTGCCTCTATATCTttCTTTTACGGCTGTCTTTGCCTGGAA  AAAACGGTGGAGGGCGTAAGTTAGTttCCACTCAACTTTAACCCG  GTCCCTGCCTCTATATCTttCCATAGTTGCTGGATTGGATATGTC  TTTTAATAGAATTTTCGATTTGCAGttCCACTCAACTTTAACCCG  GTCCCTGCCTCTATATCTttTCGCAAGTCAATACTTTCAGGTTGA  CACTATTCTAACTTCGAAGTCTAACttCCACTCAACTTTAACCCG  GTCCCTGCCTCTATATCTttTTAAAGTACGTGGCACCATCATCAT  AAAATTGTTTTGCAAGTAATGATTCttCCACTCAACTTTAACCCG  GTCCCTGCCTCTATATCTttTCTTCAAGAAAAGCTTTTGAAAATG  TTACAAATTAGCTGTGTACTCTCAGttCCACTCAACTTTAACCCG  GTCCCTGCCTCTATATCTttTATTACCGCCTGAAATTCTTTAAAA  TCCGTACCATTCGATGGGCATATCTttCCACTCAACTTTAACCCG  GTCCCTGCCTCTATATCTttCCCAAGTCGCTCAAAGTGTTGTGCC  TTTTCGAACCATTCCTGGTACTCCGttCCACTCAACTTTAACCCG  GTCCCTGCCTCTATATCTttACTCTAGTGCTTCCTCAACATTACA  AAGAATAAAATATCATGACATTTAAttCCACTCAACTTTAACCCG  GTCCCTGCCTCTATATCTttATAAGTCAGGCAACAATCTATACAC  TTTTGACATTATATCATATATTCCTttCCACTCAACTTTAACCCG  GTCCCTGCCTCTATATCTttGCATTTAGTCCTTGAAGAACTTCAA  TGTTGAACATCCACTTGATCCAGTTttCCACTCAACTTTAACCCG  GTCCCTGCCTCTATATCTttAATATGGGAGCATATATAAGCTCTA  TTGTCTCCCCATTCCATAAAATTTCttCCACTCAACTTTAACCCG | B3 |
